# Spinophilin limits metabotropic glutamate receptor 5 scaffolding to the postsynaptic density and cell type-specifically mediates excessive grooming

**DOI:** 10.1101/2022.05.24.493240

**Authors:** Cameron W. Morris, Darryl S. Watkins, Taylor Pennington, Emma H. Doud, Guihong Qi, Amber L. Mosley, Brady K. Atwood, Anthony J. Baucum

**Author notes:** To whom correspondence should be sent.

## Abstract

**Background:** Constitutive knockout of the obsessive-compulsive disorder-associated protein, disks large associated protein 3 (SAPAP3), results in repetitive motor dysfunction, such as excessive grooming, caused by increased metabotropic glutamate receptor 5 (mGluR5) activity in striatal direct- and indirect pathway medium spiny neurons (dMSNs and iMSNs, respectively). However, MSN subtype-specific signaling mechanisms that mediate mGluR5-dependent adaptations underlying excessive grooming are not fully understood. Here, we investigate the MSN subtype-specific roles of the striatal signaling hub protein, spinophilin, in mediating repetitive motor dysfunction associated with mGluR5 function.

**Methods:** Quantitative proteomics and immunoblotting were utilized to identify how spinophilin impacts mGluR5 phosphorylation and protein interaction changes. Plasticity and repetitive motor dysfunction associated with mGluR5 action was measured using our novel conditional spinophilin mouse model that had spinophilin knocked out from striatal dMSNs or/and iMSNs.

**Results:** Loss of spinophilin only in iMSNs decreased performance of a novel motor repertoire, but loss of spinophilin in either MSN subtype abrogated striatal plasticity associated with mGluR5 function and prevented excessive grooming caused by SAPAP3 knockout mice and treatment with the mGluR5-specific positive allosteric modulator (VU0360172) without impacting locomotion-relevant behavior. Biochemically, we determined spinophilin’s protein interaction correlates with grooming behavior and loss of spinophilin shifts mGluR5 interactions from lipid-raft associated proteins toward postsynaptic density (PSD) proteins implicated in psychiatric disorders.

**Conclusions:** These results identify spinophilin as a novel striatal signaling hub molecule in MSNs that cell subtype-specifically mediates behavioral, functional, and molecular adaptations associated with repetitive motor dysfunction in psychiatric disorders.

## Introduction

The sensorimotor striatum, a major basal ganglia input nucleus, integrates excitatory and modulatory inputs from diverse cortical and subcortical structures to promote the learning and execution of complex tasks [1, 2]. Perturbations within the sensorimotor striatum, or the rodent dorsolateral striatum (DLS), are associated with repetitive motor dysfunction in numerous psychiatric disorders, including obsessive-compulsive spectrum disorders (OCSDs) [3-12]. Cell type-specific adaptations in striatal direct- and indirect-pathway medium spiny neurons (dMSNs and iMSNs, respectively) within the DLS underlie repetitive and habitual actions [13-19]. Functionally, dMSNs, which express D1-type dopamine receptors, promote action execution by increasing thalamic neuronal firing rates, which in turn increase glutamatergic tone in the cortex; whereas, iMSNs, which express D2-type dopamine receptors, inhibit or temper competing motor programs by promoting the inhibition of thalamic output to the cortex, thus decreasing glutamatergic drive [20-25]. Despite bidirectional actions on basal ganglia output, dMSNs and iMSNs work in concert to integrate glutamatergic and dopaminergic signaling to promote complex motor programs, and signaling molecule perturbations within MSNs can increase the propensity to repetitively execute previously learned motor sequences, a core motor phenotype associated with OCSDs [1, 26-36].

Dysfunction in metabotropic glutamate receptor 5 (mGluR5) signaling and/or its interaction with postsynaptic density (PSD) scaffolding proteins is associated with repetitive motor dysfunction in numerous preclinical models for understanding psychiatric disorders [19, 37-42]. Of these, mutations in the striatal-enriched mGluR5 scaffold protein, disks large-associated protein 3 (SAPAP3), are associated with repetitive grooming and washing symptoms in OCSDs [43-45]. Genetic deletion of SAPAP3 in rodent results in striatal circuit abnormalities and increased mGluR5 function that promotes excessive grooming [13, 46-52], a complex sequential motor program that becomes excessively initiated and sustained despite negative consequences [53].

Reversible phosphorylation of mGluR5’s intracellular C-terminal domain is a negative feedback mechanism that promotes receptor desensitization [54-56]. Protein phosphatases, such as protein phosphatase 1 (PP1), can reverse this endocytic feedback mechanism to stabilize mGluR5 on the membrane surface [57]. Promiscuous phosphatases require targeting proteins to shuttle them into contact with their targets [58]. However, the role(s) phosphatase targeting proteins play in promoting increased mGluR5 function to mediate repetitive motor dysfunction are unknown.

Spinophilin is a striatal PSD signaling hub molecule that targets PP1 to diverse substrates [59-64]. Spinophilin promotes plasticity and motor behaviors associated with DLS function, stabilizes mGluR5 expression in the neuronal membrane, and prevents G-protein coupled receptor (GPCR) desensitization [65-74]. Recently, we determined that spinophilin interacts with SAPAP3 in mouse striatum, and overexpression of a glutamate binding deficient mGluR5 construct increased the spinophilin-SAPAP3 protein interaction [67]. However, how endogenous spinophilin mediates striatal plasticity and mGluR5 and SAPAP3-dependent repetitive motor output is unknown. Here, using a novel conditional spinophilin knockout mouse line, combined with behavioral, functional, biochemical, and proteomic approaches, we implicate spinophilin as a hub striatal signaling molecule in the striatum that mediates MSN subtype-specific adaptations underlying repetitive motor output associated with increased mGluR5 function.

## Materials and Methods

Refer to **supplement** for complete report of materials and methods.

### Animals

Animals were purchased from Jackson Laboratories or created by the university of Michigan Transgenic Animal Model Core from ES cells generated by the EUCOMMTOOLS group and obtained from the EuMMCR. Animals were maintained on a 12-hour light dark cycle (7AM:7PM) and all animal procedures were performed on 7-16 week-old male and female mice between 8AM and 5PM in accordance with the School of Science and School of Medicine Institutional Animal Care and Use Committees at IUPUI (Protocol #s SC270R, SC310R, 21090).

### Animal Behavior

Rotarod was performed as previously described [66]. All locomotion and repetitive self-grooming experiments were performed in the open field (OF) of Noldus Phenotyper Cages using a validated AI approach. Male and female mice were used for all behavior experiments, but sex was not considered as a biological variable in the present study. However, data was visualized by sex (**Supplemental Figures 18-22**).

### Electrophysiology

Mouse brain slices (350µm) containing the dorsal striatum were made from 7-10 week-old male and female mice using a Leica VT1200S vibratome, as previously described [75]. Slices recovered in artificial cerebral spinal fluid (aCSF) solution (30°C) for 1-hour then transferred to room temperature until field excitatory postsynaptic currents (fEPSPs) were recorded as described in [76] and **supplement**.

### Tissue Homogenization

Striatal tissue (whole striatum or 2 mm striatal punches from 0.5 mm coronal sections) was homogenized in low-ionic lysis buffer as previously described [64, 67, 77, 78]. Prepared lysates were used for protein concentration assay (Thermo BCA Assay, 23227 or 23252), immunoprecipitation (see below), and/or preparation of input sample by diluting lysate 1:4 in 4X Laemmli sample buffer for immunoblot analysis.

### Immunoprecipitations and Immunoblotting

Immunoprecipitation of spinophilin (Thermo; PA5-48102) or mGluR5 (Millipore; AB5675) and immunoblots were performed as previously described [64, 67, 77, 78]. Immunoblots were imaged using Odyssey CLX or Odyssey M where fluorescence intensity measurements were made using Image Studio or Empiria, respectively (LI-COR Biosciences, Lincoln, NE).

### Proteomics

mGluR5 IPs from Spino^+/+^ and Spino^-/-^ were submitted to the Indiana University Center for Proteome Analysis at the Indiana University School of Medicine for sample preparation, mass spectrometry analysis, bioinformatics, and data evaluation for the GelC-MS run and both quantitative proteomics runs similar to previously published protocols (see supplemental information for full protocols) [77, 79].

### Statistics

All statistical analyses are described fully in supplement and all t-, F-, and r-statistics from analyses are fully reported in **Table S6**.

## Results

### Creation and validation of a conditional spinophilin knockout line

Spinophilin is expressed in diverse brain regions and cell types [59, 80-82]. To directly probe spinophilin’s MSN-specific functions, we created a conditional spinophilin knockout line (**Supplemental methods, Figure S1A-B**). We biochemically validated our conditional spinophilin line (Spino^Fl/Fl^) by crossing them with a global, tamoxifen-inducible Cre-recombinase mouse line (CagCreER) and injecting these mice with tamoxifen to confirm a significant protein depletion (∼95%) in the striatum, hippocampus, cortex, and cerebellum (**Figure S1C-R**).

### Spinophilin has MSN-specific roles in mediating motor function and striatal plasticity

After confirming Cre expression depletes spinophilin protein levels, we bred Spino^Fl/Fl^ mice with Drd1- or Adora2a-Cre lines to deplete spinophilin from dMSNs (Spino^ΔdMSN^) or iMSNs (Spino^ΔiMSN^), respectively. Spinophilin protein expression was significantly reduced (∼25%) in spino^ΔdMSN^ and spino^ΔiMSN^ mice, however, PP1 levels were unaffected (**Figure 1A-E**).

**Figure 1:**
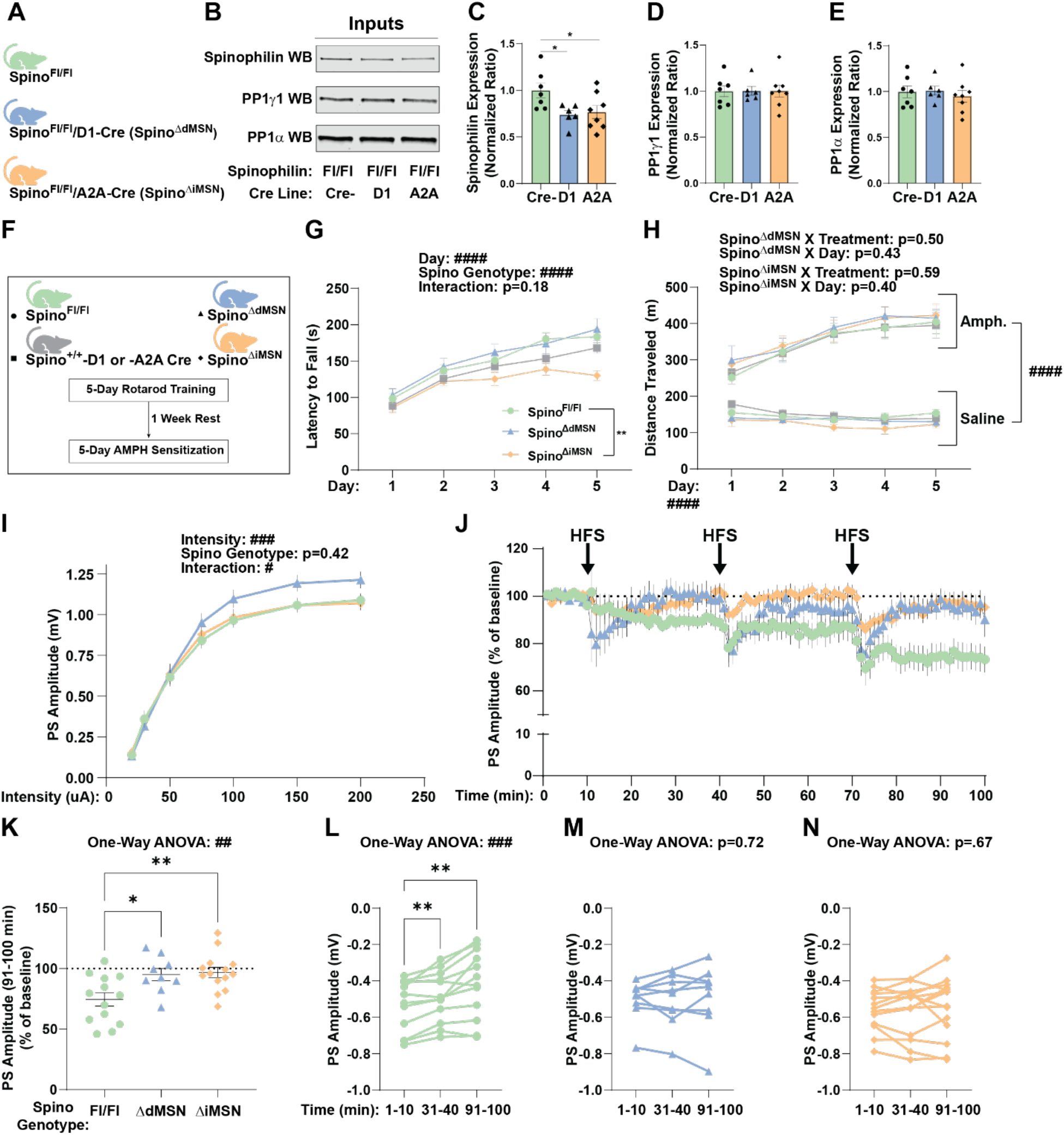
Spinophilin has MSN-specific roles in mediating motor function and striatal plasticity. **A)** Conditional spinophilin mice (Spino^Fl/Fl^) were crossed with Cre constitutively expressed under the Drd1 (D1)- or Adora2a (A2A)-promotor to deplete spinophilin expression in dMSNs (Spino^ΔdMSN^) or iMSNs (Spino^ΔiMSN^), respectively. At 2-4 months of age, 2mm striatal punches were taken from coronal slices isolated from male or female mice for **B)** immunoblot analysis of spinophilin and PP1 protein expression. One-way ANOVA with post-hoc Dunnett’s multiple comparisons test detected a significant decrease in **C)** spinophilin expression in Spino^ΔdMSN^ and Spino^ΔiMSN^ compared to control (p=0.032 and p=0.041, respectively). There was no change in **D)** PP1γ1 (p=0.99) or **E)** PP1 (p=0.75) expression. N=8 Spino^Fl/Fl^ (6 male), 6 Spino^ΔdMSN^ (2 male), 7 Spino^ΔiMSN^ (4 male). **F)** 7-9 week-old control and MSN-specific spinophilin KO mice were trained on accelerating rotarod for 5-days. After one week of rest, these same animals were treated with a sensitizing regimen of d-amphetamine (3 mg/kg) for 5-days. **G)** Two-way ANOVA with repeated measures and post-hoc Dunnett’s multiple comparisons test determined Spino^ΔiMSN^ mice performed significantly worse than Spino^Fl/Fl^ (p=0.002). N=17 Spino^Fl/Fl^ (7 male), 17 Spino^+/+^-D1 or -A2A Cre (11 male), 15 Spino^ΔdMSN^ (10 male), 15 Spino^ΔiMSN^ (5 male). **H)** Three-way ANOVAs with repeated measures detected significant day (p<0.0001) and treatment (p<0.0001) effects, but not Spino^ΔdMSN^ or Spino^ΔiMSN^ genotype or interaction effects on locomotion following 5-daily treatments with saline-or amphetamine. N=8/15 saline/AMPH-treated Spino^Fl/Fl^ (2/5 male), 9/8 saline/AMPH-treated Spino^+/+^-D1 or -A2A Cre (7/4 male), 6/9 saline/AMPH-treated Spino^ΔdMSN^ (5/5 male), 6/9 saline/AMPH-treated Spino^ΔiMSN^ (2/3 male). Field population spike amplitudes from the DLS were measured in response to **I)** stimulation intensity increases or **J)** high-frequency stimulations that result in long-term depression (LTD). Two-way ANOVA with repeated measures detected a spinophilin genotype X stimulation intensity interaction (p=0.02), however, post-hoc Šídák’s multiple comparisons test did not detect group differences at any specific intensity **(I)**. N=34 slice from 20 Spino^Fl/Fl^ mice (11 male mice), 27 slice from 17 Spino^ΔdMSN^ mice (10 male mice), 28 slices from 17 Spino^ΔiMSN^ mice (10 male mice). One-way ANOVA with post-hoc Dunnett’s multiple comparisons test determined that **K)** LTD is decreased in both Spino^ΔdMSN^ and Spino^ΔiMSN^ compared to control from 91-100 minutes (p=0.018 and p=0.004, respectively). One-way ANOVAs with repeated measures confirmed LTD in **L)** control at 31-40 and 91-100-minutes relative to the 1-10 minute baseline period (p=0.007 and p=0.0013, respectively), however, neither Spino^ΔdMSN^ nor Spino^ΔiMSN^ underwent LTD from 31-40 minutes (p=0.97 and p=0.99, respectively) or 91-100 minutes (p=0.81 and p=0.75, respectively). N=13 slice from 11 Spino^Fl/Fl^ mice (5 male mice), 9 slice from 7 Spino^ΔdMSN^ mice (3 male mice), 14 slice from 10 Spino^ΔiMSN^ mice (5 male mice). Data ± SEM. Significant two- or one-way ANOVA effects denoted by #p≤0.05, ##p≤0.01, ###p≤0.001, ####p<0.0001. Significant post-hoc tests denoted by *p≤0.05, **p≤0.01, ***p≤0.001.

We next characterized motor function in Spino^ΔdMSN^ and Spino^ΔiMSN^ mice by challenging these genotypes with an accelerating rotarod task and amphetamine-induced locomotor sensitization (**Figure 1F**), behaviors that whole-body spinophilin knockout (Spino^-/-^) decreases [66-68]. Spino^ΔiMSN^, but not Spino^ΔdMSN^, decreased rotarod performance in the later stages of this motor task (**Figure 1G**), such that Spino^ΔiMSN^ did not increase performance after day 3 (**Figure S2**). However, both Spino^ΔdMSN^ and Spino^ΔiMSN^ mice displayed acute hyperlocomotion and locomotor sensitization to repeated doses of amphetamine treatment (**Figure 1H**). These data, suggest cell type-specific actions of spinophilin mediate striatal-dependent motor actions.

To measure spinophilin’s MSN subtype-specific roles in regulating DLS network excitability and long-term plasticity we recorded field population spike amplitude responses to stimulation and long-term synaptic depression (LTD), a form of striatal plasticity that requires increased mGluR5 and D2-dopamine receptor (D_2_R) function [83, 84]. Specifically, we recorded population spike amplitudes evoked from increasing electrical stimulation intensities (input/output) or high-frequency stimulations that induce LTD (HFS-LTD) in coronal sections from control, Spino^ΔdMSN^, and Spino^ΔiMSN^ mice. We detected a spino genotype X intensity interaction suggesting Spino^ΔdMSN^ increases DLS network responses, however, we did not detect any post-hoc genotype differences within intensity groups (**Figure 1I**). Both Spino^ΔdMSN^ and Spino^ΔiMSN^ had significantly decreased HFS-LTD compared to Spino^Fl/Fl^ control. Unlike control, the population spike amplitude did not decrease in Spino^ΔdMSN^ or Spino^ΔiMSN^ following either single or multiple bouts of high-frequency stimulation (**Figure 1J-N**).

### Spinophilin dMSN- and iMSN-KO decrease excessive grooming in SAPAP3 KO mice

Excessive grooming in SAPAP3 KO mice is associated with increased mGluR5 signaling and plasticity in the DLS [13, 46]. Given that spinophilin interacts with SAPAP3 in mouse striatum, mGluR5 expression increases the spinophilin-SAPAP3 protein interaction, and MSN subtype-specific spinophilin knockout decreases DLS plasticity, we generated double knockout mice (SAPAP3 WT and KO/spinophilin MSN subtype-specific KO) to determine if spinophilin in specific MSN subtypes impacts grooming dysfunction. At 8-weeks of age, grooming and locomotion was measured in Noldus Phenotyper cages for 1-hour using a validated artificial intelligence approach (**Figure 2A-C**). Constitutive knockout of SAPAP3 significantly increased grooming duration (percent grooming) in spinophilin replete and Spino^ΔdMSN^ mice compared to their genotype controls, whereas the increased grooming was abrograted in Spino^ΔiMSN^ mice compared to their genotype controls. However, percent grooming was significantly decreased in both SAPAP3 KO/Spino^ΔdMSN^ and SAPAP3 KO/Spino^ΔiMSN^ mice compared to SAPAP3 KO/spinophilin replete mice (**Figure 2D**). This was a spinophilin-specific effect as D1-or A2A-Cre expression alone does not decrease excessive grooming in SAPAP3 KO mice (**Figure S3**).

**Figure 2:**
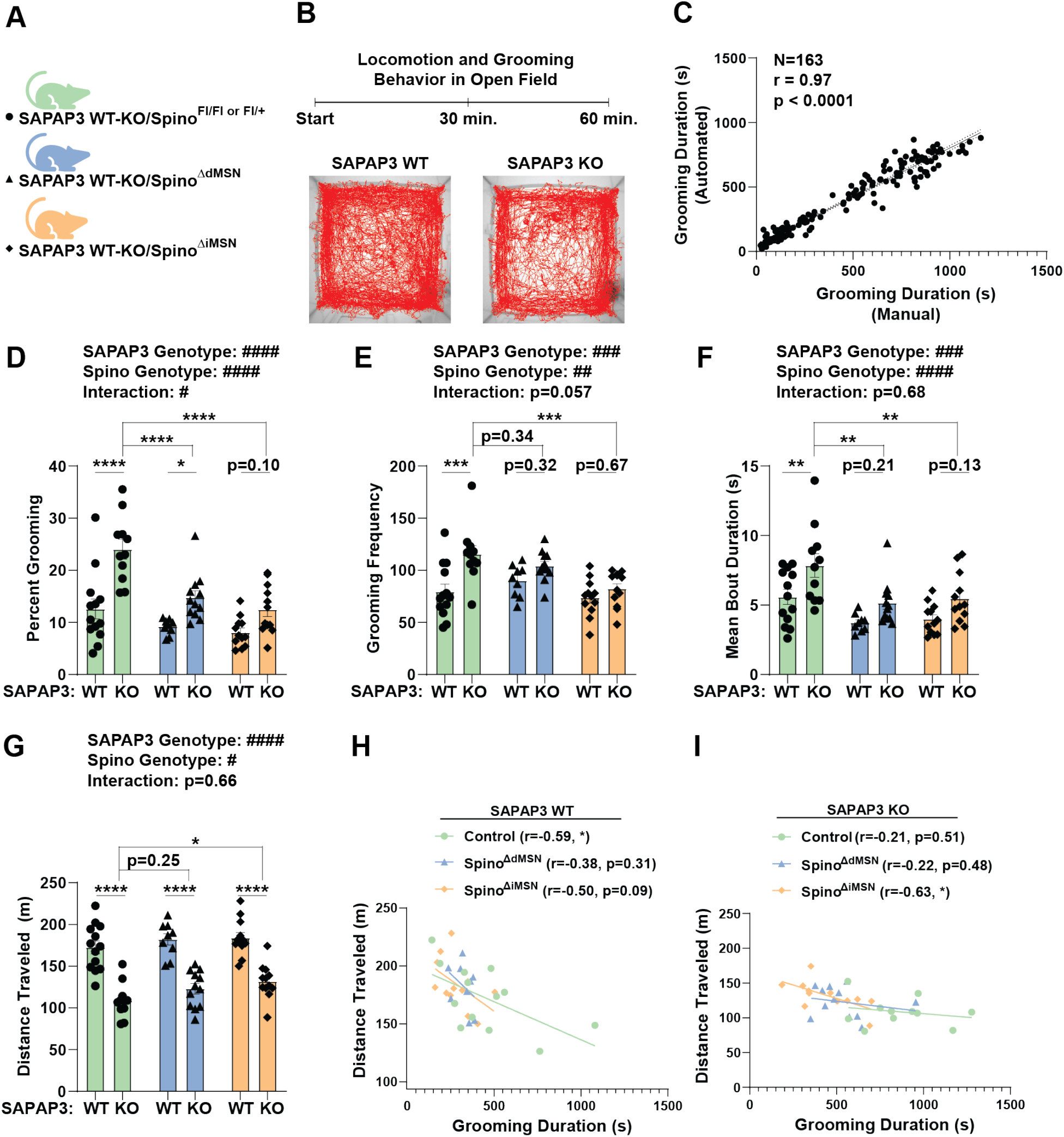
MSN-specific spinophilin knockout decreases excessive grooming in SAPAP3 deficient mice. **A)** Double knockout mice (SAPAP3 WT and KO/spinophilin MSN subtype-specific KO) were **B)** placed in the open field (OF) Noldus Phenotyper Cages for 60 minutes at 8-weeks of age to measure locomotion and grooming behavior. **C)** Grooming behavior was scored using Noldus’ behavior recognition (NBR) algorithm, which was validated by a Pearson’s correlation analysis comparing grooming duration scored manually versus the NBR algorithm (N=163 30-minute OF recordings, r=0.97, p<0.0001). Two-way ANOVAs with post-hoc Šídák’s multiple comparisons tests determined: **D)** Percent grooming was significantly decreased in SAPAP3 KO/Spino^ΔdMSN^ (p<0.0001) and SAPAP3 KO/Spino^ΔiMSN^ mice (p<0.0001) compared to SAPAP3 KO control, and a significant SAPAP3 genotype effect was detected in control (p<0.0001) and Spino^ΔdMSN^ (0.039) but not Spino^ΔiMSN^ mice (p=0.10). **E)** Grooming frequency was significantly decreased in SAPAP3 KO/Spino^ΔiMSN^ compared to SAPAP3 KO control (p=0.0005), but SAPAP3 KO/Spino^ΔdMSN^ did not affect grooming frequency relative to SAPAP3 KO control (0.34). A significant SAPAP3 genotype effect was detected in control (p=0.0002), but not Spino^ΔdMSN^ (0.32) or Spino^ΔiMSN^ mice (p=0.67). **F)** Mean grooming bout duration was significantly decreased in SAPAP3 KO/Spino^ΔdMSN^ (p=0.0014) and SAPAP3 KO/Spino^ΔiMSN^ mice (p<0.004) compared to SAPAP3 KO control. A significant SAPAP3 genotype effect was detected in control (p=0.007), but not Spino^ΔdMSN^ (p=0.21) or Spino^ΔiMSN^ mice (p=0.13). **G)** Significant SAPAP3 genotype effects were detected in control (p<0.0001), Spino^ΔdMSN^ (p<0.0001), and Spino^ΔiMSN^ mice (p<0.0001). Distance traveled was significantly increased in SAPAP3 KO/Spino^ΔiMSN^ compared to SAPAP3 KO control (p=0.032), but SAPAP3 KO/Spino^ΔdMSN^ was not different than SAPAP3 KO control (p=0.25). Grooming duration (s) and distance traveled (m) were correlated for **H)** SAPAP3 WT and **I)** SAPAP3 KO groups to determine that grooming behavior negatively correlates with locomotion in SAPAP3 WT control and SAPAP3 KO/Spino^ΔiMSN^. N=11-13 SAPAP3 WT-KO/Spino^Fl/Fl or Fl/+^ (7-5 male), N=9-12 SAPAP3 WT-KO/Spino^ΔdMSN^ (4-6 male), and N=12-12 SAPAP3 WT-KO/Spino^ΔiMSN^ (4-6 male). Data ± SEM. Significant two-ANOVA effects denoted by #p≤0.05, ##p≤0.01, ###p≤0.001, ####p<0.0001. Significant post-hoc tests denoted by *p≤0.05, **p≤0.01, ***p≤0.001, ****p<0.0001.

Grooming duration can be broken into the number of grooming bouts and the mean duration of each grooming bout. We detected SAPAP3 genotype and spinophilin genotype effects on both grooming frequency and mean grooming bout duration, such that grooming frequency was significantly reduced only in SAPAP3 KO/Spino^ΔiMSN^ and mean bout duration was significantly reduced in both SAPAP3 KO/Spino^ΔdMSN^ and SAPAP3 KO/Spino^ΔiMSN^ mice (**Figure 2E-F**), suggesting Spino^ΔdMSN^ and Spino^ΔiMSN^ mice may decrease SAPAP3 KO grooming via unique mechanisms.

In addition to grooming dysfunction, we also detected SAPAP3 genotype effects causing hypolocomotion in control, Spino^ΔdMSN^, and Spino^ΔiMSN^ mice. SAPAP3 KO/Spino^ΔiMSN^ mice had small, but significantly increased locomotion compared to SAPAP3 KO control mice (**Figure 2G-H**). Given that grooming and locomotion are competing behaviors in the OF, we correlated distance traveled and grooming duration for each genotype, but we only detected a significant negative correlation in SAPAP3 WT and SAPAP3 KO/Spino^ΔiMSN^ (**Figure 2I**). Grooming and locomotion data were also analyzed in 30-minute bins to confirm genotype differences are consistent between the first and second halves of recording (**Figure S4**).

### The interaction of spinophilin with mGluR5 is increased in SAPAP3 KO striatum and correlates with grooming duration

Given that increased striatal mGluR5 signaling is associated with grooming dysfunction in SAPAP3 KO mice we measured spinophilin’s interaction with mGluR5 in SAPAP3 KO striatum. In addition, we measured spinophilin’s interaction with the D_2_R [85]—another striatal GPCR known to decrease rodent grooming [86]. We harvested striata from 4-6 month-old SAPAP3 WT and KO mice following measurement of grooming behavior and measured the expression and interaction of spinophilin with these GPCRs as well as PP1γ1 (**Figure 3A-B**). Quantitative immunoblot analysis of striatal input samples determined that SAPAP3 KO did not affect spinophilin, mGluR5, or D_2_R expression; however, PP1γ1 expression was significantly increased in SAPAP3 KO mice (**Figure 3C-F**). Although we found no change in spinophilin’s interaction with PP1γ1, there was significantly more mGluR5 and D_2_R in spinophilin IPs from SAPAP3 KO striatum (**Figure 3G-I**). We did not detect any PP1γ1, mGluR5, or D_2_R co-IP in spinophilin IPs from spino^-/-^ striatal tissue (**Figure 3A**, lane 3), indicating co-IPs are specific. Moreover, we detected a significant Pearson’s correlation between percent grooming and the log_2_ fold-change in the spinophilin-mGluR5 or spinophilin-D_2_R interaction (**Figure 3J-K**) as well as a significant correlation between the spinophilin-mGluR5 and spinophilin-D_2_R interaction (**Figure 3L**).

**Figure 3:**
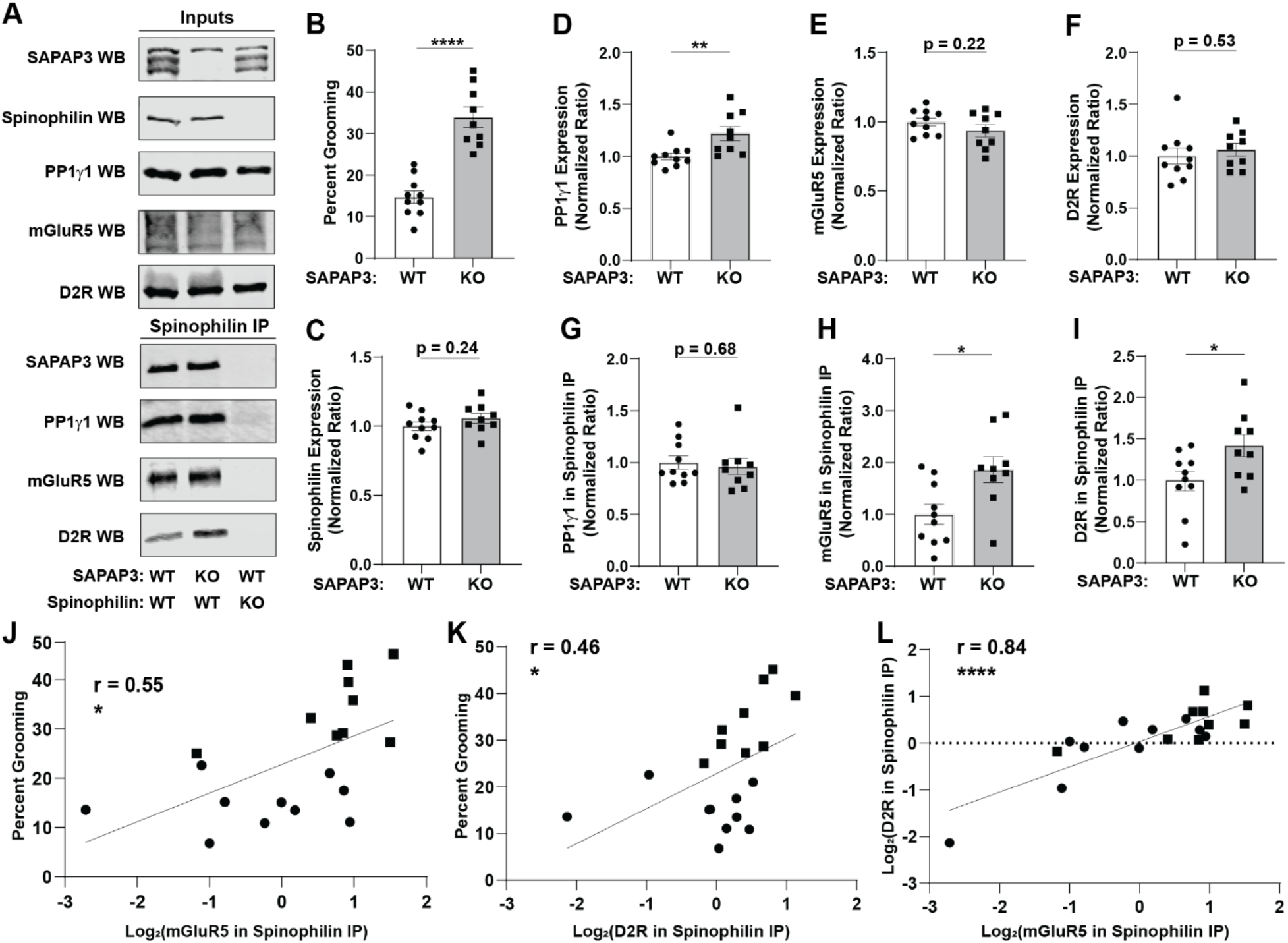
Spinophilin’s interactions with mGluR5 and D_2_R correlate with grooming behavior. **A-B)** Grooming behavior was measured in 4-6 month-old SAPAP3 WT and KO mice for 2.5 hours (p<0.0001), then striatal inputs and spinophilin immunoprecipitates (IPs) were prepared for immunoblot analysis of SAPAP3, spinophilin, PP1γ1, mGluR5 and D_2_R. Spinophilin KO striatum (lane 3) was used as a qualitative negative control to confirm the specificity of co-immunoprecipitates (co-IPs). Individual unpaired t-tests determined SAPAP3 KO did not change total protein expression of **C)** spinophilin (p=0.24), **E)** mGluR5 (p=0.22), or **F)**. D_2_R (p=0.53), however, a significant increase in **D)** PP1γ1 (p=0.007) was determined. Analysis of IPs determined significantly more **H)** mGluR5 (p=0.012) and **I)** D_2_R (p=0.033) interacting with spinophilin in SAPAP3 KO striatum, however, there is no change in spinophilin’s interaction with **G)** PP1γ1 (p=0.68). Pearson’s correlation analysis determined grooming duration positively correlates with spinophilin’s protein interaction with **J)** mGluR5 (r=0.556, p=0.013) and **K)** D_2_R (r=0.463, p=0.045), and the **L)** spinophilin-mGluR5 interaction positively correlates with the spinophilin-D_2_R interaction (r=0.840, p < 0.0001). N=10 SAPAP3 WT (7 male) and 9 SAPAP3 KO (4 male). Data ± SEM. *p≤0.05, **p≤0.01, ****p<0.0001.

### Loss of spinophilin modulates mGluR5 phosphorylation

To probe consequences of loss of spinophilin on mGluR5, we measured mGluR5 phosphorylation in spinophilin replete (Spino^+/+^) and Spino^-/-^ striatum. First, we performed striatal mGluR5 IPs from one Spino^+/+^ and Spino^-/-^ mouse and excised the mGluR5 band on a Coomassie gel for a targeted GelC-MS run to validate that we can ratiometrically quantify phosphorylation sites on mGluR5 (**Figures S5-8**). Preliminarily, we determined that loss of spinophilin increased mGluR5 phosphorylation at Ser860 and Ser870 (**Table S1**). To follow up on these preliminary results, we pooled sequential mGluR5 IPs from spino^+/+^ and spino^-/-^ striatal lysates (N=3 per genotype). These sequential IPs immunodepleted mGluR5 by ∼80%, and we found no genotype difference in immunodepletion or mGluR5 expression (**Figure 4A-C, Figure S9**). mGluR5 complexes were submitted for protein identification and ratiometric quantification using tandem mass tag-liquid chromatography/mass spectrometry (TMT-LC/MS). Given the inherent variability of IPs coupled with TMT-LC/MS [87], we utilized a Log_2_ Abundance Ratio (KO/WT) < -0.2 or > 0.2 combined with one-tailed t-tests (α<0.10) to identify decreased or increased phosphopeptides, respectively. Interestingly, loss of spinophilin not only increased the abundance of mGluR5 Ser860 and 1016 phosphorylation, but we also detected increased phosphorylation of SHANK3 Ser781 and SAPAP2 Ser983, proteins with strong genetic associations with autism spectrum disorders (ASDs) and OCSDs [88-94] (**Figure 4D**).

**Figure 4:**
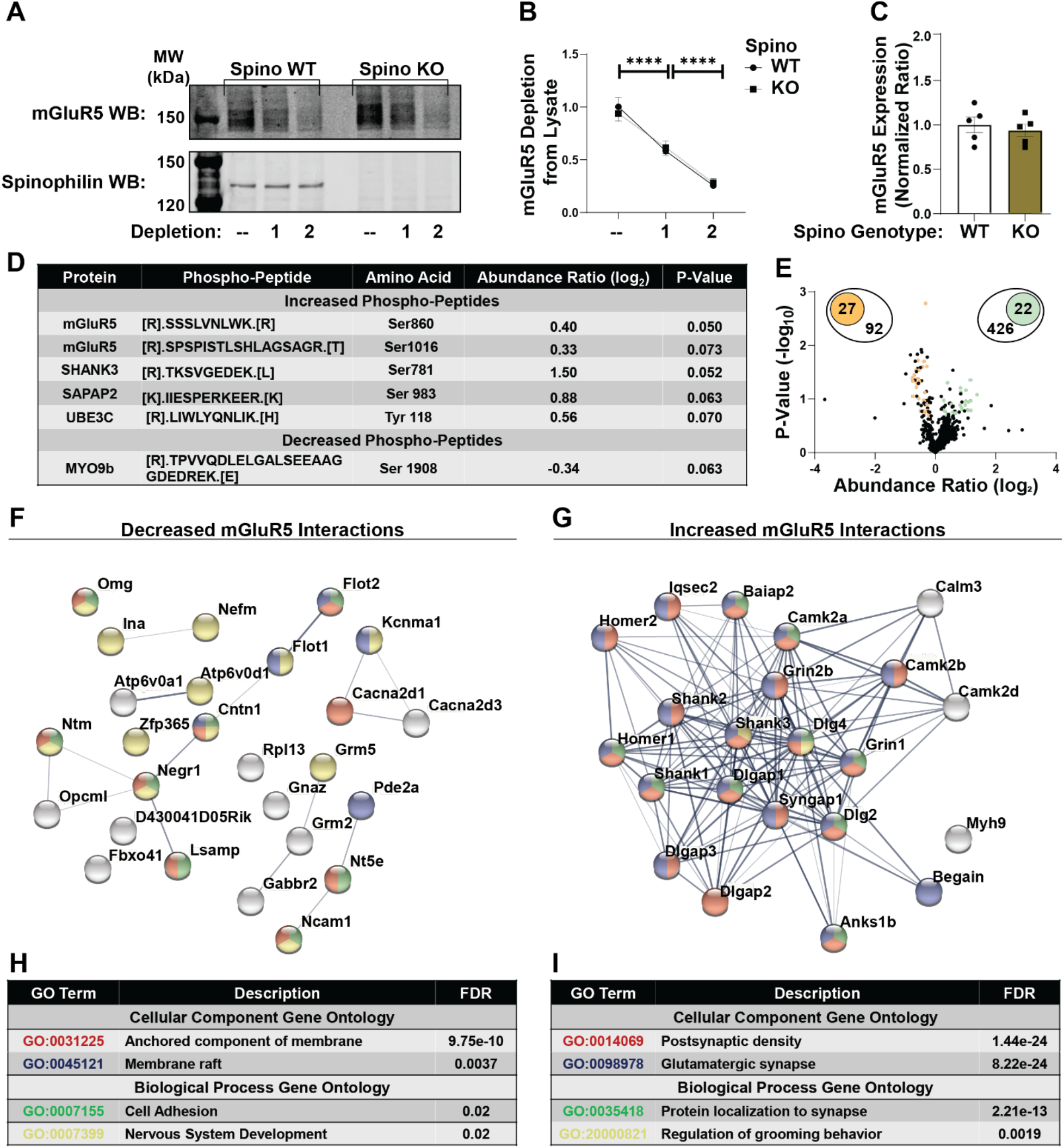
Loss of spinophilin shifts mGluR5 interactions from lipid raft assemblies toward PSD proteins implicated in psychiatric disorders. **A)** Representative sequential mGluR5 IP from spinophilin WT (Spino^+/+^) and KO (Spino^-/-^) striatal lysates. We detected no difference in **B)** mGluR5 immunodepletion or **C)** mGluR5 protein expression in spino WT and KO inputs. mGluR5 IPs were submitted to the IUSM proteomics core for TMT-MS/MS analysis. **D)** Table showing increased and decreased phospho-peptides that have Abundance Ratio (Log_2_) < -0.2 or >0.2 and one-tailed t-test p-value < 0.10. **E)** Volcano plot showing 27 decreased (orange) and 22 increased (green) interactors having Abundance Ratio (Log_2_) < -0.2 or >0.2, one-tailed t-test p-value < 0.10, and at least 6 unique peptides matching assigned protein. Protein-protein interaction (PPI) networks corresponding to the **F)** 27 decreased or **G)** 22 increased proteins were graphed in STRING. Graph edges correspond to proteins that participate in a function complex and edge boldness corresponds to the confidence of the interaction. Node colors correspond to gene ontology (GO) terms identified through function enrichment analysis of the **H)** decreased or **I)** increased PPI networks (false discovery rate (FDR) < 0.05). N=3 spino^+/+^ (3 male) and 3 spino^-/-^ (2 male).). Data ± SEM. ****p<0.0001.

### Loss of spinophilin shifts mGluR5 interactions from lipid raft assemblies toward PSD scaffolding proteins implicated in psychiatric disorders

We next analyzed protein abundance of mGluR5 co-IPs in the TMT-LC/MS dataset, which can give insight into consequences of loss of spinophilin on mGluR5 function. We identified 92 downregulated (log_2_-fold change < -0.2) and 426 upregulated (log_2_-fold change > 0.2) mGluR5 co-IPs isolated from spino^-/-^ striatum that resulted in expansive protein-protein interaction (PPI) networks (**Figures S10-11**). We refined this list of interactors by filtering for significantly decreased/increased proteins (p<0.10 from one-tailed t-tests) that matched at least 6 unique peptides. We identified 27 decreased and 22 increased high-confidence mGluR5 interactors in spino^-/-^ striatum (orange and green points in **Figure 4E**, respectively). We graphed these high-confidence interactors in STRING to generate decreased and increased PPI networks that were used for functional enrichment analysis (**Figure 4F-I, Table S3-4**). We determined that spinophilin knockout decreased mGluR5 interactions with glycosylphosphatidylinositol (GPI)-anchored proteins, a class of proteins localized to specialized microdomains in the plasma membrane known as lipid rafts [95], and increased mGluR5 interactions with numerous PSD scaffolding proteins, including SAPAP3 and SHANK3. Shifting mGluR5 interactions toward PSD proteins in spino^-/-^ striatum was associated with multiple Biological Process GO terms, one of which was “regulation of grooming behavior”.

We validated these protein interaction findings by submitting a second, small cohort of mGluR5 IPs (N=2 per genotype) (**Table S5**) for TMT-LC/MS. We replicated that loss of spinophilin decreased (log_2_-fold change < 0.2) mGluR5 protein interactions with GPI-anchored proteins and increased (log_2_-fold change > 0.2) mGluR5 interactions with PSD proteins implicated in psychiatric disorders, including SAPAP3 and SHANK3. *Spinophilin MSN-specifically Decreases Grooming Caused by the mGluR5 positive allosteric modulator (PAM), VU0360172 (VU’172)*.

To directly determine if spinophilin mediates excessive grooming by regulating mGluR5 function we pharmacologically increased grooming behavior by treating mice with the mGluR5 PAM, VU’172, that selectively increases grooming in wild type, but not mGluR5 KO mice [13]. Specifically, we measured grooming behavior in control, Spino^ΔdMSN^, and Spino^ΔiMSN^ mice for 30-minutes before and after an I.P. injection of vehicle or VU’172 (20 mg/kg). We also measured the grooming response to VU’172 in Spino^-/-^ mice to determine any additive or antagonistic effects when spinophilin is knocked out of both MSN subtypes (**Figure 5A**). We found no treatment group effects on percent grooming or distance traveled in the pre-injection period (**Figure S12**); however, percent grooming was decreased, and distance traveled was increased in Spino^-/-^ compared to control mice, a published phenotype of Spino^-/-^ mice [96]. Vehicle treatment increased grooming (a grooming phenotype that is also decreased in mGluR5 KO mice); however grooming was further increased by VU’172 treatment in Spino^Fl/+ or Fl/Fl^, Spino^-/-^, and Spino^+/+^-D1 or -A2A Cre control mice (**Figure 5B, S13**). Furthermore, we detected significant treatment X spinophilin genotype interaction on percent grooming, such that we did not detect a VU’172 treatment effect on percent grooming in Spino^ΔdMSN^ and Spino^ΔiMSN^ mice, and percent grooming in VU’172-treated Spino^ΔdMSN^ and Spino^ΔiMSN^ mice was not significantly different from vehicle-treated control mice (**Figure 5B**). Post-injection percent grooming data were also analyzed in 5-minute bins to confirm genotype effects were consistent across 30-minute recording (**Figure S14**). Collectively, these data suggest MSN subtype-specific spinophilin knockout, but not global knockout, abrogates VU’172-induced grooming.

**Figure 5:**
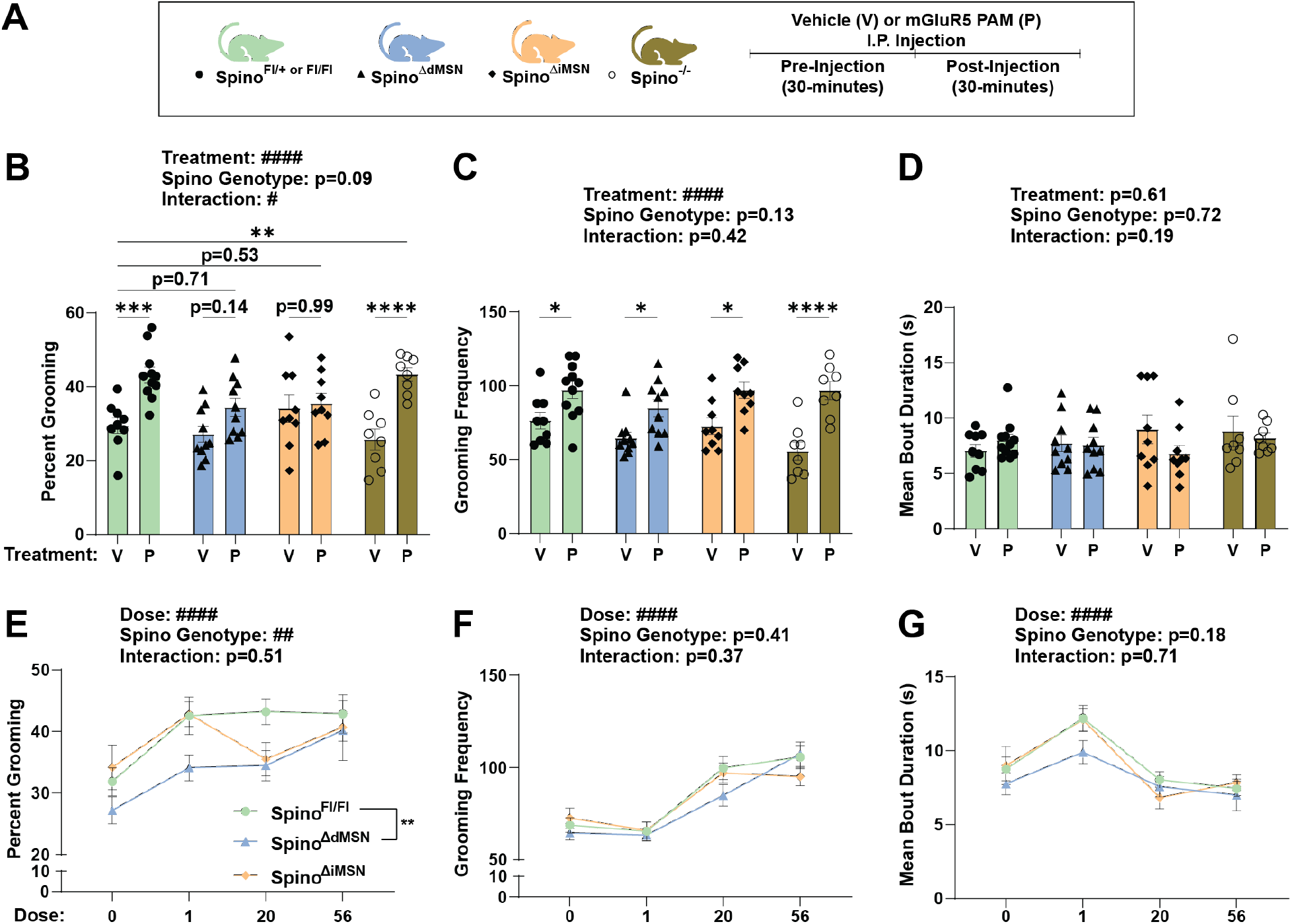
Spinophilin MSN-specifically decreases grooming caused by the mGluR5 PAM, VU0360172. **A)** Control and MSN-specific spinophilin knockout mice were placed in Noldus Phenotyper Cages and basal behavior (pre-injection) was measured for 30-minutes. Following a pre-injection period, mice were removed from the arena and given an I.P. injection of vehicle (V) or the mGluR5 PAM (P), VU0360172 (VU’172) (20mg/kg). Animals were placed back into the same pre-injection arena immediately following the I.P. injection and behavior was recorded for an additional 30-minutes (post-injection). Two-way ANOVAs with post-hoc Šídák’s multiple comparisons tests determined a significant VU’172 treatment effect on **B)** percent grooming in control (p=0.0007) and Spino^-/-^ (p<0.0001), but not Spino^Δ.dMSN^ (p=0.139) or Spino^Δ.iMSN^ (p=0.99) mice. Furthermore, percent grooming in the VU’172-treated Spino^Δ.dMSN^ and Spino^Δ.iMSN^ groups was not different from control vehicle (p=0.71 and p=0.53, respectively), whereas Spino^-/-^ was significantly increased from control vehicle (p=0.002). **C)** VU’172 (20 mg/kg) increased grooming frequency in control (p=0.03), Spino^Δ.dMSN^ (p=0.03), Spino^Δ.iMSN^ (p=0.01), and Spino^-/-^ (p<0.0001). **D)** VU’172 (20 mg/kg) or spinophilin genotype did not affect mean grooming bout duration. N=9V/10P Spino^Fl/+ or Fl/Fl^ (3/3 male), 10V/10P spino^Δ.dMSN^ (5/6 male), and 9V/9P spino^Δ.iMSN^ (4/4 male), 8V/8P spino^-/-^ (3/4 male). **E-G)** Separate cohorts of Spino^Fl/Fl^ control, Spino^Δ.dMSN^, and Spino^Δ.iMSN^ mice were treated with vehicle, 1, or 56 mg/kg VU’172 and combined with existing 20 mg/kg data (B-D) to generate 4-point dose response curves. Two-way ANOVAs with post-hoc Dunnett’s multiple comparisons tests determined **E)** percent grooming was significantly decreased in Spino^Δ.dMSN^ relative to Spino^Fl/Fl^ control (p=0.004), but Spino^Δ.iMSN^ was not different (0.56). However, spinophilin genotype did not significantly affect **F)** grooming frequency or **G)** mean grooming bout duration. N=11-22 Spino^Fl/Fl or Fl/+^ (8-14 male), 8-11 Spino^Δ.dMSN^ (4-6 male), and 9-11 Spino^Δ iMSN^ (2-5 male). Data ± SEM. Significant two-ANOVA effects denoted by #p≤0.05 and ####p<0.0001. Significant post-hoc tests denoted by *p≤0.05, **p≤0.01, ***p≤0.001 ****p<0.0001.

Although we detected a significant treatment effect on grooming frequency, we found no genotype or genotype X treatment interaction (**Figure 5C**). Also, neither VU’172 (20 mg/kg) nor spinophilin genotype affected mean grooming bout duration (**Figure 5D**). Alternatively, we detected a significant treatment, genotype, and interaction effect on locomotion in OF, but post-hoc tests determined this is due to increased locomotion in vehicle-treated Spino^-/-^ mice. Grooming duration in VU’172-treated control negatively correlated with distance traveled; however, we did not detect significant correlations in the other groups (**Figure S15**).

To determine how loss of spinophilin in both MSN subtypes modifies VU’172-induced grooming, we bred Spino^ΔdMSN^ and Spino^ΔiMSN^ mice to knockout spinophilin from both MSN subtypes (Spino^ΔPanMSN^). We treated this genotype with VU’172 (20 mg/kg) and compared to the aforementioned Spino^Fl/Fl or Fl/+^ and Spino^-/-^ VU’172 grooming response. In contrast to the individual cell type knockouts, there was no significant effect of the double knockout compared to the control or Spino^-/-^ groups in percent grooming, grooming frequency, and mean bout duration across the 3 groups (**Figure S16**), suggesting that the lack of effect on VU’172-induced grooming observed in Spino^-/-^ mice was due to loss of spinophilin in both MSN subtypes having an antagonistic effect on VU’172-induced grooming.

To further understand these unexpected findings, we performed a VU’172 dose response curve in Spino^Fl/Fl^ control mice using a dose range consistent with published studies using VU’172 in mice [97-99]. We not only found significant VU’172 treatment effects on percent grooming but determined percent grooming at low doses is driven by increased mean grooming bout duration, whereas, at high doses, percent grooming is driven by increased grooming frequency (**Figure S17**). Given this, we treated separate cohorts of Spino^ΔdMSN^ and Spino^ΔiMSN^ with a low (1 mg/kg) and high (56 mg/kg) dose of VU’172 and combined these data with our existing 20 mg/kg data (**Figure 5B-D**) to create a 4-point dose response curve for grooming duration, grooming frequency, and mean grooming bout duration (**Figure E-G**). We determined that Spino^ΔdMSN^ decreased percent grooming compared to control. Although the two-way ANOVA did not detect spinophilin genotype effects or interactions on grooming frequency or mean grooming bout duration, we noticed a subtle, non-significant, decrease in Spino^ΔdMSN^ mean grooming bout duration at 1 mg/kg and grooming frequency at 20 mg/kg, suggesting Spino^ΔdMSN^-dependent grooming decreases are not exclusively driven by duration or frequency alone.

## Discussion

Conditional spinophilin knockout from dMSNs or iMSNs decreased excessive grooming caused by constitutive knockout of the striatal-enriched PSD scaffold, SAPAP3. Excessive grooming in SAPAP3 KO mice is associated with MSN subtype-specific adaptations that increase dMSN function relative to neighboring iMSNs in the DLS [13]. Both of these striatal abnormalities—excessive grooming and increased dMSN function— are decreased by treatment with the mGluR5 negative allosteric modulator (NAM), MTEP [13]. We determined that spinophilin’s protein interaction with mGluR5 is increased in SAPAP3 KO mice, and that MSN subtype-specific spinophilin knockout also prevents increased grooming duration caused by the mGluR5-specific PAM, VU’172. Collectively, these data suggest that spinophilin expression in striatal MSNs mediates mGluR5-dependent excessive grooming.

Given that MSN subtype-specific spinophilin knockout prevented excessive grooming in SAPAP3 KO mice and VU’172-treated mice (20 mg/kg), it was unexpected that whole-body loss of spinophilin failed to decrease VU’172 grooming. We further confirmed this by detecting a strong VU’172 grooming response despite depleting spinophilin from both MSN subtypes (Spino^ΔPanMSN^), suggesting spinophilin mediates mGluR5-dependent excessive grooming in an antagonistic, MSN subtype-specific manner. We further probed this surprising effect by comparing 4-point VU’172 dose response curve (DRC) from control, Spino^ΔdMSN^, and Spino^ΔdMSN^ mice. Surprisingly, we found unique DRCs across these genotypes, such that Spino^ΔdMSN^ significantly decreased the percent grooming DRC, whereas the Spino^ΔiMSN^ effect on percent grooming was limited to the 20 mg/kg dose.

Increased grooming duration (percent grooming) is achieved, at least in part, by increased grooming bout initiation (grooming frequency) or by sustained grooming bouts (mean grooming bout duration). Increased dMSN function is essential for initiating and sustaining complex motor programs, including rodent self-grooming [16, 86, 100]. Interestingly, SAPAP3 KO/Spino^ΔdMSN^ mice had decreased mean grooming duration compared to controls, suggesting Spino^ΔdMSN^ may be important in sustaining complex motor programs. Although Spino^ΔdMSN^ decreased the VU’172 percent grooming DRC, this was not solely due to decreasing grooming frequency or mean grooming bout duration. Rather, we only saw subtle decreases in both grooming frequency and mean grooming bout duration, suggesting Spino^ΔdMSN^ likely integrates unique aspects of grooming behavior to promote increased grooming duration in response to VU’172.

Decreasing iMSN function with a D_2_R agonist decreases grooming initiation, duration, and sequence completion [86, 100]. Interestingly, spino^ΔiMSN^ mice had decreased grooming frequency and mean grooming bout duration in SAPAP3 KO mice. Furthermore, spinophilin’s interaction with D_2_R was increased in SAPAP3 KO striatum and correlated with grooming behavior. In addition to mediating excessive grooming, spinophilin expression in iMSNs was also necessary for increased performance of a novel repetitive motor sequence needed to master the accelerating rotarod task, another motor behavior associated with D_2_R function [101]. Collectively, these data suggest spinophilin expression in iMSNs may be critical for executing and completing complex sequential motor programs.

Recently, Tecuapetla and colleagues determined that optogenetic inhibition of striatal iMSNs in the dorsomedial striatum (DMS) decreased excessive grooming in SAPAP3 KO mice [102]. While we have identified that MSN subtype-specific loss of spinophilin decreases excessive grooming, it is unclear if spinophilin functions in specific striatal subregions to mediate excessive grooming. Furthermore, while the D1- and A2A-Cre lines utilized herein are highly expressed within striatal dMSNs or iMSNs, it is possible that spinophilin mediates excessive grooming by functioning in cell types outside the striatum through residual extra-striatal Cre expression or in a non-cell-autonomous manner. However, both Spino^ΔdMSN^ and Spino^ΔiMSN^ mice had abrogated HFS-LTD in the DLS—plasticity that requires mGluR5 and D_2_R function [83, 84]. The role of decreased LTD limiting pathological grooming is consistent with studies showing that SAPAP3 KO mice undergo *increased* short-term plasticity due to increased mGluR5 function in the DLS [46]. Furthermore, the increased DLS network responses in Spino^ΔdMSN^ mice is also an opposite phenotype of preclinical excessive grooming models [52, 103]. Future studies will build upon the foundation laid herein to delineate spinophilin’s cell-autonomous and non-autonomous functions in specific striatal subregions in mediating plasticity associated with repetitive motor dysfunction.

Spinophilin knockout-derived primary cortical neurons have increased mGluR5-dependent MAPK signaling and intracellular calcium mobilization ([Ca^2+^]) that leads to increased mGluR5 endocytosis [69]. Loss of spinophilin significantly upregulated striatal mGluR5 phosphorylation at Ser860 and Ser1016, a protein kinase A (PKA) consensus site and CaMKII site, respectively [104]. mGluR5 phosphorylation was also upregulated (Abundance Ratio (Log2) > 0.2) at Ser839 and Ser908, protein kinase C (PKC) sites [105, 106]. Although the function of mGluR5 Ser860 phosphorylation is unknown, phosphorylation of Ser1016, 839, and 908 promote increased MAPK signaling, [Ca^2+^]_i_ mobilization, and mGluR5 desensitization [54, 55, 104-106]. These data further implicate spinophilin as a regulator of striatal mGluR5 signaling. Future studies will elucidate how phosphorylation of these mGluR5 sites govern its signaling, interactions, and localization within the postsynaptic membrane.

Excessive grooming in both SAPAP3 KO and SHANK3 complete KO mice— preclinical models for understanding OCSDs and ASDs, respectively—is decreased by mGluR5 NAMs. Grooming dysfunction in both these preclinical models is associated with decreased mGluR5 scaffolding to PSD proteins [13, 19]. Strikingly, in addition to MSN subtype-specific spinophilin knockout decreasing mGluR5-depenent excessive grooming, loss of spinophilin in the striatum increased mGluR5 interactions with PSD scaffolding proteins implicated in psychiatric disorders, including SAPAP3 and SHANK3, and decreased mGluR5 interactions with lipid raft-associated membrane proteins. Not only is lipid raft dysfunction associated with psychiatric disorders like the ASD, fragile X syndrome (FXS) [42, 107-109], mGluR5 can differentially signal depending on its interactions with lipid raft signaling complexes or PSD scaffolding proteins [110-112]. While the signaling pathways underlying mGluR5-depenent excessive grooming have not been identified, we hypothesize that our MSN subtype-specific spinophilin knockout mice can elucidate which mGluR5 signaling pathway(s) cause excessive grooming, results that may hold broad therapeutic potential across mGluR5-opathies.

In addition to MSN subtype-specific spinophilin KO decreasing two models of excessive grooming, we also identified that spino^ΔiMSN^ plateaus the performance of a novel repetitive motor sequence using an accelerating rotarod task, but neither spino^ΔdMSN^ nor spino^ΔiMSN^ affected locomotor sensitization to psychostimulant. Therefore, we hypothesize that MSN subtype-specific spinophilin knockout selectively disrupts complex sequential motor programs without impacting overall locomotor output, such as basal, hypo- or hyperlocomotion. Due to this, we postulate that our novel MSN subtype-specific spinophilin knockout models are a critical tool to elucidate unique cell autonomous and/or non-autonomous signaling pathways underlying the initiation and/or sustainment of sequential motor programs.

## Acknowledgements

Mass spectrometry work performed herein was done by the Indiana University School of Medicine Center for Proteome Analysis. Acquisition of the IUSM Proteomics instrumentation used for this project was provided by the Indiana University Precision Health Initiative. The proteomics work was supported, in part, by the Indiana Clinical and Translational Sciences Institute funded, in part by Award Number UL1TR002529 from the National Institutes of Health, National Center for Advancing Translational Sciences, Clinical and Translational Sciences Award and the Cancer Center Support Grant for the IU Simon Comprehensive Cancer Center (Award Number P30CA082709) from the National Cancer Institute. We acknowledge and thank Wanda Filipiak & Galina Gavrilina for embryo injections for the initial generation of Spino^Fl/Fl^ mice as well as the entire excellent Transgenic Animal Model Core (in particular, Anna LaForest, Elizabeth Hughes, Corey Ziebell, and Dr. Thomas Saunders) and the University of Michigan’s Biomedical Research Core Facilities for their generation of these mice. Funding for the generation of these mice and completion of studies comes from an R21/R33 award from the National Institutes of Drug Abuse (R21/R33 DA041876 to AJB), Department of Biology/School of Science at IUPUI, Department of Pharmacology and Toxicology Startup Funds, Strategic Research Initiative Funds (Indiana University School of Medicine and Stark Neurosciences Research Institute). We appreciate the feedback from all members of the Baucum laboratory over the years on this project.

## Data Availability

Proteomics datasets from GelC-MS, TMT-LC/MS run 1, and TMT-LC/MS run 2 experiments were uploaded to ProteomeXchange (accession number pending).

## References

1. Graybiel, A.M., Habits, rituals, and the evaluative brain. Annu Rev Neurosci, 2008. 31: p. 359–87.

2. Jahanshahi, M., et al., A fronto–striato–subthalamic–pallidal network for goal-directed and habitual inhibition. Nature Reviews Neuroscience, 2015. 16: p. 719.

3. Abbott, A.E., et al., Repetitive behaviors in autism are linked to imbalance of corticostriatal connectivity: a functional connectivity MRI study. Soc Cogn Affect Neurosci, 2018. 13(1): p. 32–42.

4. Anticevic, A., et al., Global resting-state functional magnetic resonance imaging analysis identifies frontal cortex, striatal, and cerebellar dysconnectivity in obsessive-compulsive disorder. Biol Psychiatry, 2014. 75(8): p. 595–605.

5. Beucke, J.C., et al., Abnormally high degree connectivity of the orbitofrontal cortex in obsessive-compulsive disorder. JAMA Psychiatry, 2013. 70(6): p. 619–29.

6. O’Sullivan, R.L., et al., Reduced basal ganglia volumes in trichotillomania measured via morphometric magnetic resonance imaging. Biological Psychiatry, 1997. 42(1): p. 39–45.

7. Sakai, Y., et al., Corticostriatal functional connectivity in non-medicated patients with obsessive-compulsive disorder. Eur Psychiatry, 2011. 26(7): p. 463–9.

8. Atmaca, M., et al., Volumetric MRI study of key brain regions implicated in obsessive-compulsive disorder. Prog Neuropsychopharmacol Biol Psychiatry, 2007. 31(1): p. 46–52.

9. Pujol, J., et al., Mapping structural brain alterations in obsessive-compulsive disorder. Arch Gen Psychiatry, 2004. 61(7): p. 720–30.

10. Harrison, B.J., et al., Brain corticostriatal systems and the major clinical symptom dimensions of obsessive-compulsive disorder. Biol Psychiatry, 2013. 73(4): p. 321–8.

11. Di Martino, A., et al., Aberrant striatal functional connectivity in children with autism. Biological psychiatry, 2011. 69(9): p. 847–856.

12. Grossberg, S. and D. Kishnan, Neural Dynamics of Autistic Repetitive Behaviors and Fragile X Syndrome: Basal Ganglia Movement Gating and mGluR-Modulated Adaptively Timed Learning. Front Psychol, 2018. 9: p. 269.

13. Ade, K.K., et al., Increased Metabotropic Glutamate Receptor 5 Signaling Underlies Obsessive-Compulsive Disorder-like Behavioral and Striatal Circuit Abnormalities in Mice. Biol Psychiatry, 2016. 80(7): p. 522–33.

14. O’Hare, J.K., et al., Pathway-Specific Striatal Substrates for Habitual Behavior. Neuron, 2016. 89(3): p. 472–9.

15. Rothwell, P.E., et al., Autism-associated neuroligin-3 mutations commonly impair striatal circuits to boost repetitive behaviors. Cell, 2014. 158(1): p. 198–212.

16. Rothwell, P.E., et al., Input-and output-specific regulation of serial order performance by corticostriatal circuits. Neuron, 2015. 88(2): p. 345–356.

17. Sheng, M.J., et al., Emergence of stable striatal D1R and D2R neuronal ensembles with distinct firing sequence during motor learning. Proc Natl Acad Sci U S A, 2019.

18. Wang, W., et al., Striatopallidal dysfunction underlies repetitive behavior in Shank3-deficient model of autism. J Clin Invest, 2017. 127(5): p. 1978–1990.

19. Wang, X., et al., Altered mGluR5-Homer scaffolds and corticostriatal connectivity in a Shank3 complete knockout model of autism. Nature Communications, 2016. 7(1): p. 11459.

20. Gerfen, C.R. and D.J. Surmeier, Modulation of striatal projection systems by dopamine. Annual review of neuroscience, 2011. 34: p. 441–466.

21. Cui, G., et al., Concurrent activation of striatal direct and indirect pathways during action initiation. Nature, 2013. 494(7436): p. 238–42.

22. Tecuapetla, F., et al., Balanced activity in basal ganglia projection pathways is critical for contraversive movements. Nat Commun, 2014. 5: p. 4315.

23. Trusel, M., et al., Coordinated Regulation of Synaptic Plasticity at Striatopallidal and Striatonigral Neurons Orchestrates Motor Control. Cell Rep, 2015. 13(7): p. 1353–1365.

24. Corbit, L.H., J.L. Muir, and B.W. Balleine, Lesions of mediodorsal thalamus and anterior thalamic nuclei produce dissociable effects on instrumental conditioning in rats. Eur J Neurosci, 2003. 18(5): p. 1286–94.

25. Gerfen, C.R., et al., D1 and D2 dopamine receptor-regulated gene expression of striatonigral and striatopallidal neurons. Science, 1990. 250(4986): p. 1429–1432.

26. Greengard, P., The neurobiology of slow synaptic transmission. Science, 2001. 294(5544): p. 1024–30.

27. Uematsu, K., et al., Protein kinase A directly phosphorylates metabotropic glutamate receptor 5 to modulate its function. J Neurochem, 2015. 132(6): p. 677–86.

28. Voulalas, P.J., et al., Metabotropic glutamate receptors and dopamine receptors cooperate to enhance extracellular signal-regulated kinase phosphorylation in striatal neurons. J Neurosci, 2005. 25(15): p. 3763–73.

29. Francesconi, A. and R.M. Duvoisin, Opposing effects of protein kinase C and protein kinase A on metabotropic glutamate receptor signaling: selective desensitization of the inositol trisphosphate/Ca2+ pathway by phosphorylation of the receptor-G protein-coupling domain. Proc Natl Acad Sci U S A, 2000. 97(11): p. 6185–90.

30. Martiros, N., A.A. Burgess, and A.M. Graybiel, Inversely Active Striatal Projection Neurons and Interneurons Selectively Delimit Useful Behavioral Sequences. Curr Biol, 2018. 28(4): p. 560–573.e5.

31. Barnes, T.D., et al., Activity of striatal neurons reflects dynamic encoding and recoding of procedural memories. Nature, 2005. 437(7062): p. 1158–61.

32. Jin, X. and R.M. Costa, Start/stop signals emerge in nigrostriatal circuits during sequence learning. Nature, 2010. 466(7305): p. 457–62.

33. Jin, X., F. Tecuapetla, and R.M. Costa, Basal ganglia subcircuits distinctively encode the parsing and concatenation of action sequences. Nat Neurosci, 2014. 17(3): p. 423–30.

34. Jog, M.S., et al., Building neural representations of habits. Science, 1999. 286(5445): p. 1745–9.

35. Gillan, C.M., et al., Counterfactual processing of economic action-outcome alternatives in obsessive-compulsive disorder: further evidence of impaired goal-directed behavior. Biol Psychiatry, 2014. 75(8): p. 639–46.

36. Gillan, C.M., et al., Enhanced avoidance habits in obsessive-compulsive disorder. Biol Psychiatry, 2014. 75(8): p. 631–8.

37. D’Antoni, S., et al., Dysregulation of group-I metabotropic glutamate (mGlu) receptor mediated signalling in disorders associated with Intellectual Disability and Autism. Neurosci Biobehav Rev, 2014. 46 Pt 2: p. 228–41.

38. Matosin, N., et al., Shifting towards a model of mGluR5 dysregulation in schizophrenia: consequences for future schizophrenia treatment. Neuropharmacology, 2017. 115: p. 73–91.

39. Mehta, M.V., M.J. Gandal, and S.J. Siegel, mGluR5-antagonist mediated reversal of elevated stereotyped, repetitive behaviors in the VPA model of autism. PLoS One, 2011. 6(10): p. e26077.

40. Pop, A.S., et al., Fragile X syndrome: a preclinical review on metabotropic glutamate receptor 5 (mGluR5) antagonists and drug development. Psychopharmacology (Berl), 2014. 231(6): p. 1217–26.

41. Stewart, S.E., et al., Genome-wide association study of obsessive-compulsive disorder. Molecular Psychiatry, 2013. 18(7): p. 788–798.

42. Ronesi, J.A., et al., Disrupted Homer scaffolds mediate abnormal mGluR5 function in a mouse model of fragile X syndrome. Nat Neurosci, 2012. 15(3): p. 431–40, s1.

43. Bienvenu, O.J., et al., Sapap3 and pathological grooming in humans: Results from the OCD collaborative genetics study. Am J Med Genet B Neuropsychiatr Genet, 2009. 150b(5): p. 710–20.

44. Naaz, S., et al., Association of SAPAP3 allelic variants with symptom dimensions and pharmacological treatment response in obsessive-compulsive disorder. Exp Clin Psychopharmacol, 2020.

45. Zuchner, S., et al., Multiple rare SAPAP3 missense variants in trichotillomania and OCD. Mol Psychiatry, 2009. 14(1): p. 6–9.

46. Chen, M., et al., Sapap3 deletion anomalously activates short-term endocannabinoid-mediated synaptic plasticity. J Neurosci, 2011. 31(26): p. 9563–73.

47. Corbit, V.L., et al., Strengthened Inputs from Secondary Motor Cortex to Striatum in a Mouse Model of Compulsive Behavior. J Neurosci, 2019. 39(15): p. 2965–2975.

48. Hadjas, L.C., C. Lüscher, and L.D. Simmler, Aberrant habit formation in the Sapap3-knockout mouse model of obsessive-compulsive disorder. Sci Rep, 2019. 9(1): p. 12061.

49. Hadjas, L.C., et al., Projection-specific deficits in synaptic transmission in adult Sapap3-knockout mice. Neuropsychopharmacology, 2020.

50. Wan, Y., et al., Circuit-selective striatal synaptic dysfunction in the Sapap3 knockout mouse model of obsessive-compulsive disorder. Biol Psychiatry, 2014. 75(8): p. 623–30.

51. Wan, Y., G. Feng, and N. Calakos, Sapap3 deletion causes mGluR5-dependent silencing of AMPAR synapses. J Neurosci, 2011. 31(46): p. 16685–91.

52. Welch, J.M., et al., Cortico-striatal synaptic defects and OCD-like behaviours in Sapap3-mutant mice. Nature, 2007. 448(7156): p. 894–900.

53. Kalueff, A.V., et al., Neurobiology of rodent self-grooming and its value for translational neuroscience. Nat Rev Neurosci, 2016. 17(1): p. 45–59.

54. Gereau, R.W.t. and S.F. Heinemann, Role of protein kinase C phosphorylation in rapid desensitization of metabotropic glutamate receptor 5. Neuron, 1998. 20(1): p. 143–51.

55. Ko, S.J., et al., PKC phosphorylation regulates mGluR5 trafficking by enhancing binding of Siah-1A. J Neurosci, 2012. 32(46): p. 16391–401.

56. Alagarsamy, S., S.D. Sorensen, and P.J. Conn, Coordinate regulation of metabotropic glutamate receptors. Curr Opin Neurobiol, 2001. 11(3): p. 357–62.

57. Kliewer, A., R.K. Reinscheid, and S. Schulz, Emerging Paradigms of G Protein-Coupled Receptor Dephosphorylation. Trends Pharmacol Sci, 2017. 38(7): p. 621–636.

58. Cohen, P.T., Protein phosphatase 1--targeted in many directions. J Cell Sci, 2002. 115(Pt 2): p. 241–56.

59. Allen, P.B., C.C. Ouimet, and P. Greengard, Spinophilin, a novel protein phosphatase 1 binding protein localized to dendritic spines. Proc Natl Acad Sci U S A, 1997. 94(18): p. 9956–61.

60. Colbran, R.J., et al., Association of brain protein phosphatase 1 with cytoskeletal targeting/regulatory subunits. J Neurochem, 1997. 69(3): p. 920–9.

61. Ragusa, M.J., et al., Spinophilin directs protein phosphatase 1 specificity by blocking substrate binding sites. Nat Struct Mol Biol, 2010. 17(4): p. 459–64.

62. Baucum, A.J., 2nd, et al., Identification and validation of novel spinophilin-associated proteins in rodent striatum using an enhanced ex vivo shotgun proteomics approach. Mol Cell Proteomics, 2010. 9(6): p. 1243–59.

63. Sarrouilhe, D., et al., Spinophilin: from partners to functions. Biochimie, 2006. 88(9): p. 1099–113.

64. Watkins, D.S., et al., Proteomic Analysis of the Spinophilin Interactome in Rodent Striatum Following Psychostimulant Sensitization. Proteomes, 2018. 6(4).

65. Allen, P.B., et al., Distinct roles for spinophilin and neurabin in dopamine-mediated plasticity. Neuroscience, 2006. 140(3): p. 897–911.

66. Edler, M.C., et al., Mechanisms Regulating the Association of Protein Phosphatase 1 with Spinophilin and Neurabin. ACS Chem Neurosci, 2018. 9(11): p. 2701–2712.

67. Morris, C.W., et al., The association of spinophilin with disks large-associated protein 3 (SAPAP3) is regulated by metabotropic glutamate receptor (mGluR) 5. Mol Cell Neurosci, 2018. 90: p. 60–69.

68. Areal, L.B., et al., Neuronal scaffolding protein spinophilin is integral for cocaine-induced behavioral sensitization and ERK1/2 activation. Mol Brain, 2019. 12(1): p. 15.

69. Di Sebastiano, A.R., et al., Role of Spinophilin in Group I Metabotropic Glutamate Receptor Endocytosis, Signaling, and Synaptic Plasticity. J Biol Chem, 2016. 291(34): p. 17602–15.

70. Fujii, S., et al., Spinophilin inhibits the binding of RGS8 to M1-mAChR but enhances the regulatory function of RGS8. Biochem Biophys Res Commun, 2008. 377(1): p. 200–4.

71. Kurogi, M., et al., Effects of spinophilin on the function of RGS8 regulating signals from M2 and M3-mAChRs. Neuroreport, 2009. 20(13): p. 1134–9.

72. Wang, Q., et al., Spinophilin blocks arrestin actions in vitro and in vivo at G protein-coupled receptors. Science, 2004. 304(5679): p. 1940–4.

73. Wang, X., et al., Spinophilin/neurabin reciprocally regulate signaling intensity by G protein-coupled receptors. Embo j, 2007. 26(11): p. 2768–76.

74. Wang, X., et al., Spinophilin regulates Ca2+ signalling by binding the N-terminal domain of RGS2 and the third intracellular loop of G-protein-coupled receptors. Nat Cell Biol, 2005. 7(4): p. 405–11.

75. Karahan, H., et al., Deletion of Abi3 gene locus exacerbates neuropathological features of Alzheimer’s disease in a mouse model of Aβ amyloidosis. Sci Adv, 2021. 7(45): p. eabe3954.

76. Yin, H.H., et al., Dynamic reorganization of striatal circuits during the acquisition and consolidation of a skill. Nat Neurosci, 2009. 12(3): p. 333–41.

77. Salek, A.B., et al., Spinophilin regulates phosphorylation and interactions of the GluN2B subunit of the N-methyl-d-aspartate receptor. J Neurochem, 2019. 151(2): p. 185–203.

78. Hiday, A.C., et al., Mechanisms and Consequences of Dopamine Depletion-Induced Attenuation of the Spinophilin/Neurofilament Medium Interaction. Neural Plast, 2017. 2017: p. 4153076.

79. Grecco, G.G., et al., A multi-omic analysis of the dorsal striatum in an animal model of divergent genetic risk for alcohol use disorder. J Neurochem, 2021. 157(4): p. 1013–1031.

80. Muhammad, K., et al., Presynaptic spinophilin tunes neurexin signalling to control active zone architecture and function. Nat Commun, 2015. 6: p. 8362.

81. Muly, E.C., et al., Subcellular distribution of neurabin immunolabeling in primate prefrontal cortex: comparison with spinophilin. Cereb Cortex, 2004. 14(12): p. 1398–407.

82. Ouimet, C.C., et al., Cellular and subcellular distribution of spinophilin, a PP1 regulatory protein that bundles F-actin in dendritic spines. J Comp Neurol, 2004. 479(4): p. 374–88.

83. Kreitzer, A.C. and R.C. Malenka, Dopamine modulation of state-dependent endocannabinoid release and long-term depression in the striatum. Journal of Neuroscience, 2005. 25(45): p. 10537–10545.

84. Wu, Y.-W., et al., Input-and Cell-Type-Specific Endocannabinoid-Dependent LTD in the Striatum. Cell Reports, 2015. 10(1): p. 75–87.

85. Smith, F.D., G.S. Oxford, and S.L. Milgram, Association of the D2 dopamine receptor third cytoplasmic loop with spinophilin, a protein phosphatase-1-interacting protein. J Biol Chem, 1999. 274(28): p. 19894–900.

86. Berridge, K.C. and J.W. Aldridge, Super-stereotypy I: enhancement of a complex movement sequence by systemic dopamine D1 agonists. Synapse, 2000. 37(3): p. 194–204.

87. Stein, B.D., et al., Comparison of CRISPR Genomic Tagging for Affinity Purification and Endogenous Immunoprecipitation Coupled with Quantitative Mass Spectrometry To Identify the Dynamic AMPKα2 Interactome. Journal of Proteome Research, 2019. 18(10): p. 3703–3714.

88. Wu, K., et al., Glutamate system genes and brain volume alterations in pediatric obsessive-compulsive disorder: a preliminary study. Psychiatry Res, 2013. 211(3): p. 214–20.

89. Durand, C.M., et al., Mutations in the gene encoding the synaptic scaffolding protein SHANK3 are associated with autism spectrum disorders. Nature Genetics, 2007. 39(1): p. 25–27.

90. Prasad, C., et al., Genetic evaluation of pervasive developmental disorders: the terminal 22q13 deletion syndrome may represent a recognizable phenotype. Clinical Genetics, 2000. 57(2): p. 103–109.

91. Wilson, H.L., et al., Molecular characterisation of the 22q13 deletion syndrome supports the role of haploinsufficiency of <em>SHANK3/PROSAP2</em> in the major neurological symptoms. Journal of Medical Genetics, 2003. 40(8): p. 575–584.

92. Chien, W.-H., et al., Deep exon resequencing of DLGAP2 as a candidate gene of autism spectrum disorders. Molecular Autism, 2013. 4(1): p. 26.

93. Marshall, C.R., et al., Structural Variation of Chromosomes in Autism Spectrum Disorder. The American Journal of Human Genetics, 2008. 82(2): p. 477–488.

94. Pinto, D., et al., Functional impact of global rare copy number variation in autism spectrum disorders. Nature, 2010. 466(7304): p. 368–372.

95. Um, J.W. and J. Ko, Neural Glycosylphosphatidylinositol-Anchored Proteins in Synaptic Specification. Trends Cell Biol, 2017. 27(12): p. 931–945.

96. Zhang, Y., et al., Spinophilin-deficient mice are protected from diet-induced obesity and insulin resistance. Am J Physiol Endocrinol Metab, 2020. 319(2): p. E354–e362.

97. Noetzel, M.J., et al., Functional impact of allosteric agonist activity of selective positive allosteric modulators of metabotropic glutamate receptor subtype 5 in regulating central nervous system function. Mol Pharmacol, 2012. 81(2): p. 120–33.

98. Notartomaso, S., et al., Pharmacological enhancement of mGlu1 metabotropic glutamate receptors causes a prolonged symptomatic benefit in a mouse model of spinocerebellar ataxia type 1. Molecular Brain, 2013. 6(1): p. 48.

99. Zuena, A.R., et al., In Vivo Non-radioactive Assessment of mGlu5 Receptor-Activated Polyphosphoinositide Hydrolysis in Response to Systemic Administration of a Positive Allosteric Modulator. Frontiers in Pharmacology, 2018. 9.

100. Berridge, K.C. and J.W. Aldridge, Super-stereotypy II: enhancement of a complex movement sequence by intraventricular dopamine D1 agonists. Synapse, 2000. 37(3): p. 205–15.

101. Augustin, S.M., et al., Dopamine D2 receptor signaling on iMSNs is required for initiation and vigor of learned actions. Neuropsychopharmacology, 2020. 45(12): p. 2087–2097.

102. Ramírez-Armenta, K.I., et al., Optogenetic inhibition of indirect pathway neurons in the dorsomedial striatum reduces excessive grooming in Sapap3-knockout mice. Neuropsychopharmacology, 2022. 47(2): p. 477–487.

103. Peça, J., et al., Shank3 mutant mice display autistic-like behaviours and striatal dysfunction. Nature, 2011. 472(7344): p. 437–42.

104. Raka, F., et al., Ca2+/Calmodulin-dependent protein Kinase II interacts with group I Metabotropic Glutamate and facilitates Receptor Endocytosis and ERK1/2 signaling: role of β-Amyloid. Molecular Brain, 2015. 8(1): p. 21.

105. Kim, C.H., et al., Protein kinase C phosphorylation of the metabotropic glutamate receptor mGluR5 on Serine 839 regulates Ca2+ oscillations. J Biol Chem, 2005. 280(27): p. 25409–15.

106. Lee, J.H., et al., Calmodulin dynamically regulates the trafficking of the metabotropic glutamate receptor mGluR5. Proc Natl Acad Sci U S A, 2008. 105(34): p. 12575–80.

107. Kalinowska, M., C. Castillo, and A. Francesconi, Quantitative Profiling of Brain Lipid Raft Proteome in a Mouse Model of Fragile X Syndrome. PLOS ONE, 2015. 10(4): p. e0121464.

108. Toupin, A., et al., Association of lipid rafts cholesterol with clinical profile in fragile X syndrome. Scientific Reports, 2022. 12(1): p. 2936.

109. Wang, H., Lipid rafts: a signaling platform linking cholesterol metabolism to synaptic deficits in autism spectrum disorders. Front Behav Neurosci, 2014. 8: p. 104.

110. Francesconi, A., R. Kumari, and R.S. Zukin, Regulation of group I metabotropic glutamate receptor trafficking and signaling by the caveolar/lipid raft pathway. J Neurosci, 2009. 29(11): p. 3590–602.

111. Mao, L., et al., The scaffold protein Homer1b/c links metabotropic glutamate receptor 5 to extracellular signal-regulated protein kinase cascades in neurons. J Neurosci, 2005. 25(10): p. 2741–52.

112. Tu, J.C., et al., Homer binds a novel proline-rich motif and links group 1 metabotropic glutamate receptors with IP3 receptors. Neuron, 1998. 21(4): p. 717–26.

