## Supplemental methods, Table S1, and Figures for "Spinophilin limits metabotropic glutamate receptor 5 scaffolding to the postsynaptic density and cell type-specifically mediates excessive grooming"

### *Supplemental Information*

### Supplemental Methods

<u>Animals</u>

The following genotypes were obtained from Jackson Laboratories or the mutant mouse regional resource center (MMRRC): Whole-body spinophilin knockout (KO) mice (B6N(Cg)-*Ppp1r9b*$^{tm1.1(KOMP)Vlcg}$/J; bred in house after obtaining from Jackson laboratories); SAPAP3 KO mice (B6.129-Dlgap3$^{tm1Gfng}$/J); CagCreER (Tg(CAG-cre/Esr1*)5Amc; Drd1-Cre (036916-UCD B6.FVB(Cg)-Tg(Drd1a-cre)FK150Gsat/Mmucd); Adora2a-Cre (MMRRC strain 036158-UCD, B6.FVB(Cg)-Tg(Adora2a-cre)KG139Gsat/Mmucd). Conditional spinophilin KO mice were created by the University of Michigan Transgenic Animal Model Core from ES cells generated by the EUCOMMTOOLS group and obtained from the EuMMCR containing a targeted insert with a beta-gal reporter and neomycin selection cassette surrounded by FRT sites as well as loxP sites flanking exon 3 of the spinophilin gene. Upon crossing these animals with mice expressing Flp recombinase, only the loxp sites surrounding exon 3 of the spinophilin gene, Ppp1r9b, remained. Initial insert was confirmed by the EuMMCR using long-range PCR. Following Flp recombination the following DNA insert remained:
TGCCAGAGGTCCATAGGTCAGGAAGGCGCATAACGATACCACGATATCAACAAGT
TTGTACAAAAAAGCAGGCTGGCGCCGGAACCGAAGTTCCTATTCCGAAGTTCCTAT
TCCGAAGTTCCTATTCTCTAGAAAGTATAGGAACTTCGTCGAGATAACTTCGTATAG
CATACATTATACGAAGTTATGTCGAGATATCTAGACCCAGCTTTCTTGTACAAAGTG

GTTGATATCTCTATAGTCGCAGTAGGCGGCAGGGCAGGTGTAGCTAGGCAGTGGG TATCCACGTTTGGTCAGTGACGCTCTGGGCTCAGAAGAGAATAATAGGAGATCAAA GGCTCAGGGCTAAGACCTGGCTCAGTCCCCATCCCAGCCACACACCCTTATCTGT ATGCACAATAGCCTACCACACCCCAGCCACCCCTCACACACCCTGCTGCCTGTCA GCTCGTTGGATAGGGCAAAGGGCAGGAAGTAACCCGTATGAGTTAGCCTGGCAAA AGGGAGGTGGGAGGAACCATAATTCTTTCCCATCATGGGGTGACTCGTAAGCTTG GGTAGAAGATCCCTGCCAGTCATGGCTTACCCCCCTTCTGCCTCACAGACTCTGA GGGCTTGGGCATC. Genotyping primers complementary to the endogenous Ppp1r9b sequence surrounding the insert were made (forward and reverse primers, respectively: TGCCAGAGGTCCATAGGTCAGG and GATGCCCAAGCCCTCAGAGT) to amplify the wild-type band lacking insert (392 base pairs) and mutant band containing insert (624 base pairs) (**Figure S1B**). All genotypes described above were backcrossed at least 6 generations and/or maintained on C57BL/6 background. SAPAP3$^{+/-}$ and/or Spinophilin$^{Fl/Fl,\ Fl/+,\ or\ +/+}$ mice were bred with Drd1- and Adora2a-Cre lines to obtain all genotypes utilized herein. Male and female mice were used for all experiments, but sex was not considered as a biological variable in the present study. However, behavior and electrophysiology data were visualized by sex (**Figures S18-22**).

Animal Behavior

*Rotarod and Amphetamine-induced locomotor sensitization*. Male and female, 7-9 week old control (Apino$^{Fl/Fl}$ or Apino$^{+/+}$-D1 or -A2A Cre), Spino$^{\Delta dMSN}$ (Spino$^{Fl/Fl}$/D1-Cre), and Spino$^{\Delta iMSN}$ (Spino$^{Fl/Fl}$/A2A-Cre) mice were challenged for 5 days on an accelerating rotarod task (4-40 rpm in 300 s, 3 cm width, Rotamex-5, Columbus Instruments) for 3 successive trials on 5 consecutive days, similar to previously described [1]. Briefly, a series of photocell beams located above the rotating rod with a temporal resolution of 0.1 rpm (0.1 cm/sec) detected when the mouse was no longer on the rod. To circumvent any false fall recordings, any mouse that was able to grip the bar and rotate with the bar for 2 rotations or greater was considered to have failed the trial at the time of the first rotation. Latency to fall was documented for each mouse, and a 120–180 s rest was given to all mice before the next trial.

Following training on the accelerating rotarod, the same cohort of animals were given 7 days of rest and were then treated with a sensitizing regiment of d-amphetamine (3 mg/kg) similar to previously described in Morris et al. [2] with the exception that this experiment was performed in Noldus Phenotyper Cages (30 cm X 30 cm X 30 cm). Briefly, mice were given an intraperitoneal (i.p.) injection of 3.0 mg/kg of d-amphetamine (Sigma) or the saline volume equivalent to the control groups. d-Amphetamine was administered once every 24 hours for five consecutive days. Immediately after each i.p. injection, mice were placed in Noldus Phenotyper Cages and locomotion (distance traveled) was measured for 60-minutes using a single identifier localized to the center of the mouse's body.

<u>Grooming Behavior</u>

*Grooming Classification by Noldus Behavior Recognition Module.* Noldus Behavior Recognition Module is a machine learning algorithm that utilizes nose, center, and tail identifiers to measure region-specific changes in animal orientation, which is input into a probabilistic model to predict rodent behavior. We optimized the placement of the nose, center, and tail identifiers (contour-based detection settings) using age-matched C57BL/6 mice. Once optimized, we measured the accuracy of the algorithm in predicting grooming behavior, which we defined as the initiation of any phase of grooming along the cephalo-caudal axis targeting the nose, face, ears, or body [3, 4]. Grooming duration was manually scored for 163, 30-minute videos and was compared to the results obtained from Noldus' Behavior Recognition Algorithm (default probability settings) (**Figure 2C**). Pearson Correlation analysis determined the Noldus Behavior Recognition Module has a high accuracy when predicting grooming behavior (N=163, r=0.97, $p \leq 0.0001$), therefore, to increase consistency in grooming classification and prevent bias in manual scoring, we utilized this algorithm to measure changes in grooming behavior for the experiments below.

*SAPAP3 Grooming*. Singly-housed male and female, 8-week old control (SAPAP3 WT/Spino$^{Fl/Fl\ or\ Fl/+}$ or SAPAP3 KO/Spino$^{Fl/Fl\ or\ Fl/+}$ or SAPAP3 WT/Spino$^{+/+}$/D1-Cre or SAPAP3 WT/Spino$^{+/+}$/A2A-Cre), Spino$^{\Delta dMSN}$ (SAPAP3 WT/Spino$^{Fl/Fl}$/D1-Cre or SAPAP3 KO/Spino$^{Fl/Fl}$/D1-Cre), and Spino$^{\Delta iMSN}$ (SAPAP3 WT/Spino$^{Fl/Fl}$/A2A-Cre or SAPAP3 KO/Spino$^{Fl/Fl}$/A2A-Cre) mice were placed in Noldus Phenotyper Cages (30 cm x 30 cm x 30 cm) where locomotion (distance traveled and average velocity) and grooming (grooming duration, grooming frequency, mean grooming bout duration) behavior was measured for 1-hour.

*mGluR5 PAM Grooming*. Male and female 7-9 week old control (Spino$^{Fl/Fl\ or\ Fl/+}$ or Spino$^{+/+}$-D1 or -A2A Cre), Spino$^{\Delta dMSN}$ (Spino$^{Fl/Fl}$/D1-Cre), and Spino$^{\Delta iMSN}$ (Spino$^{Fl/Fl}$/A2A-Cre) mice were placed in Noldus Phenotyper Cages (30 cm x 30 cm x 30 cm) where locomotion (distance traveled and average velocity) and grooming (grooming duration, grooming frequency, mean grooming bout duration) behavior was measured for 30-

minutes (pre-injection period). Following the pre-injection period, animals were given an i.p. injection of VU0360172 (1, 3, 10, 20, 30, or 56 mg/kg) or an equivalent volume of vehicle (10% tween-80 in $H_2O$) and placed back into the same arena and behavior was measured for an additional 30 minutes (post-injection period).

Electrophysiology

Male and female 7-10 week old control (Spino$^{Fl/Fl}$), Spino$^{\Delta dMSN}$ (Spino$^{Fl/Fl}$/D1-Cre), and Spino$^{\Delta iMSN}$ (Spino$^{Fl/Fl}$/A2A-Cre) mice were isoflurane-anesthetized and the brain was rapidly dissected out. 350 μm thick coronal brain slices containing the dorsal striatum were made using a Leica VT1200S vibratome. The cutting solution was a sucrose-based ice-cold solution that contained 194 mM sucrose, 30 mM NaCl, 4.5 mM KCl, 1 mM MgCl2, 26 mM NaHCO3, 1.2 mM NaH2PO4, 10 mM glucose and was saturated with 95% O2/5% CO2. Slices were then transferred to a 95% O2/5% CO2-saturated artificial cerebral spinal fluid (aCSF) solution that contained 124 mM NaCl, 4.5 mM KCl, 1 mM MgCl2, 26 mM NaHCO3, 1.2 mM NaH2PO4, 10 mM glucose, and 2 mM CaCl2. Slices were stored at 30°C for 1-h then transferred to room temperature until recordings were performed.

Recordings were performed in 95% O2/5% CO2-saturated aCSF at 30° to 32°C with continuous perfusion at a rate of 1 to 2 ml/min. Field excitatory postsynaptic currents (fEPSPs) were recorded with micropipettes filled with 1 M NaCl using a Multiclamp 700B amplifier and Clampex software (Molecular Devices). Tungsten stereotrodes (~1 MW) were used to stimulate the dorsolateral striatum and population spike amplitudes (in mV) were measured. Stimulation parameters were adjusted using a constant current isolated stimulator (Digitimer). Using the stimulation strength that produced 50% of the maximum response, a stable baseline was recorded for 10 min stimulating at 0.05 Hz. Long-term depression (LTD) was induced similar to previous work [5]. This was done by applying two 1-s trains of 100 pulses (10 μs per pulse) delivered at 100 Hz, with 10 s between trains. Changes in population spike amplitudes were recorded for 30 min after induction monitoring at a rate of 0.05 Hz. This process, applying the LTD protocol and recording the response for 30 min, was repeated two more times. Data were expressed as a percentage of change with respect to the average baseline.

Tissue Homogenization

Striatal tissue (whole striatum or 2mm striatal punches from 0.5mm coronal sections) was homogenized in low-ionic lysis buffer (0.01 M DTT, 0.005 M EDTA, 0.002 M Tris-HCl pH 7.5, 1% Triton X-100) containing protease inhibitors (Bimake) and phosphatase inhibitors (20 mM sodium fluoride, 20 mM sodium orthovanadate, 20 mM β-glycerophosphate, and 10 mM sodium pyrophosphate; Sigma-Aldrich, St. Louis, MO or ThermoFisher). Homogenates were sonicated for 15s at 25% amplitude, rotated for 4ºC for 5 minutes, and centrifuged at 15,000 X g for 10 minutes at 4ºC. Resulting supernatants were transferred to clean microfuge tube where prepared lysates were used for protein concentration assay (Thermo BCA Assay, 23227 or 23252), immunoprecipitation (see below), and/or preparation of input sample by diluting lysate 1:4 in 4X Laemmli sample buffer for immunoblot analysis.

Immunoprecipitations

*Spinophilin*. Grooming behavior was measured in Noldus Phenotyper Cages (see below) for 2.5 hours in 4-6 month old SAPAP3 WT or KO mice. After behavior recording, whole striata were dissected, flash frozen, and stored at -80ºC. Lysates were prepared by weighing and homogenizing (as described above) in low-ionic lysis buffer (45 μL/mg of tissue) and 500 μL of striatal lysate (1.5 – 2.0 mg total protein) was added to microfuge tube and 2 μg of sheep-spinophilin primary antibody (Thermo; PA5-48102) was spiked into the lysate and tubes incubated overnight at 4ºC with rotation. The following morning, 15 μL of Protein G magnetic beads was spiked into the sample and incubated two-hours at 4ºC with rotation. Beads were washed as described above and resuspended in 40 μL of 2X Laemmli sample buffer.

*mGluR5.* Dissected whole striatum was immediately homogenized (as described above) in 1 mL of low-ionic lysis buffer and 750 μL of striatal lysate was added to a microfuge tube and 1 μg of rabbit-mGluR5 primary antibody (Millipore; AB5675) was spiked into the lysate and tubes incubated for 2.5 hours at 4ºC with rotation, then 15 μL of Protein G magnetic beads was added, and samples were incubated for an additional 2.5 hours at 4ºC with rotation. Samples were placed on magnetic tube rack and the lysate was transferred into a new microfuge tube where an aliquot of lysate was taken to measure immunodepletion and an additional 1 μg of rabbit-mGluR5 primary antibody was spiked into the lysate and samples were incubated overnight at 4ºC with rotation. The next day, another aliquot of lysates was taken to measure immunodepletion. For each round of IP, Protein G magnetic beads were washed three times with immunoprecipitation wash buffer (150 mM NaCl, 50 mM Tris-HCl pH 7.5, 0.5% (v/v) Triton X-100) and washed beads were resuspended in 30 μL of 1X PBS. Beads from the first and second IP were combined after confirming the sequential IPs sufficiently depleted mGluR5 from lysate via immunoblot (see below) analysis (**Figure S9**).

Immunoblots

Immunoblotting was performed as previously described [2, 6-8]. Briefly, inputs (12.5 μg total protein for 2mm tissue punches, 30 μg total protein for whole striatum) or immunoprecipitates were separated by SDS-PAGE and blotted using the following antibodies: sheep-spinophilin (Thermo; PA5-48102), rabbit-SAPAP3 (Millipore; ABN325), mouse-mGluR5 (Millipore; MABN540), goat-PP1γ1 (SantaCruz; sc-6108), mouse-PP1α (SantaCruz; sc-7482) and mouse-D2R (SantaCruz; sc-5303). Appropriate infrared secondary antibodies were used (Donkey anti goat, donkey anti rabbit, or donkey anti mouse conjugated to Alexa Fluor 690 or 780; ThermoFisher or Jackson ImmunoResearch) and imaged using Odyssey CLX or Odyssey M where fluorescence intensity measurements were made using Image Studio or Empiria, respectively (LI-COR Biosciences, Lincoln, NE). We previously confirmed linearity of signal intensity for spinophilin, SAPAP3, and mGluR5 [2]. At least two independent cohorts of animals were utilized for all immunoblot experiments, and each gel was run with at least an N of 2 per condition for each cohort. Samples were normalized to the average control value within each gel to allow for comparisons across gels.

Proteomics

Sample preparation, mass spectrometry analysis, bioinformatics, and data evaluation for quantitative proteomics and phosphoproteomics experiments were performed in collaboration with the Indiana University Center for Proteome Analysis at the Indiana University School of Medicine similarly to previously published protocols [9].

GelC-MS

*Sample Preparation.* Samples collected from SDS-PAGE were de-stained with 25 mM ammonium bicarbonate in 50 % acetonitrile (ACN). For all digestion steps, a volume sufficient to cover the gel pieces were used. Next, 10 mM DTT in 25 mM ammonium bicarbonate was added to reduce disulfides. 25 mM iodoacetamide was then added to alkylate free sulfhydryl groups. After addition of iodoacetamide, the reaction was incubated in the dark for 45 minutes. Gel pieces were incubated in 25 mM ammonium bicarbonate. The gel pieces were then dehydrated with 25 mM ammonium bicarbonate in 50% ACN. The samples were then placed in a rotary vacuum and centrifuged until dry and subsequently digested with 12.5 ng/μl trypsin in 25 mM ammonium bicarbonate at 37°C overnight. The supernatant was collected from all samples. The remaining gel pieces were washed with 5% formic acid in 50% ACN and were vortexed and sonicated for 5 minutes. The supernatants were collected and pooled with previous supernatants and were submitted to the Indiana University Proteomics Core Facility for analysis, where Peptides were separated on an Ultimate 3000 HPLC with loading on a 5 cm C18 trap column Acclaim™ PepMap™ 100 (3 μm particle size, 75 μm diameter; Thermo Scientific, Cat No: 164946) followed by a 15 cm PepMap RSLC C18 EASY-Spray column (Thermo Scientific, Cat No: ES900) and analyzed using a Q-Exactive Plus mass spectrometer (Thermo Fisher Scientific) operated in positive ion mode. Solvent B was increased from 5%-35% over 75 min, to 90% over 2 min, back to 3% over 2 minutes (Solvent A: 95% water, 5% acetonitrile, 0.1% formic acid; Solvent B: 100% acetonitrile, 0.1% formic acid).

A data dependent top 15 method was used with MS scan range of 200-2000 m/z, resolution of 70,000, AGC target 3e6, maximum IT of 100 ms. MS2 resolution of 17,500, fixed first mass 100 m/z, normalized collision energy of 30, isolation window of 4 m/z, target AGC of 1e5, and maximum IT of 50 ms. Dynamic exclusion of 30 sec, charge exclusion of 1, 7, 8, >8 and isotopic exclusion parameters were used.

*Data analysis.* Data were analyzed using Proteome Discoverer 2.4 (Thermo Fisher Scientific). Sequest HT search used a Uniprot *Mus musculus* database downloaded 01_09_2017, full trypsin cleavage with max of 2 missed cleavage sites, precursor tolerance of 10 ppm, fragment mass tolerance of 0.02 Da, static modifications of carbamidomethyl on C, dynamic modifications (max 3 per peptide) of oxidation on M, phosphorylation on S, T, or Y, and acetylation, met-loss or met-loss plus acetylation on protein N-termini. Fixed value PSM validator was used as the FDR node with a maximum delta Cn of 0.05. Spectra of interest were hand annotated using Xcalibur (Thermo Fisher Scientific).

<u>TMT-LC/MS</u>

*Sample Preparation.* After IP and wash steps, beads (3 WT and 3 KO in the first batch and 2 WT and 2 KO in the second) were submitted to the Center for Proteome analysis. 30 µL of 8 M Urea in 100 mM Tris pH 8.5 was added and proteins were reduced with 5 mM tris(2-carboxyethyl)phosphine hydrochloride (TCEP, Sigma-Aldrich Cat No: C4706) for 30 minutes at room temperature. The resulting free cysteine thiols were alkylated with 10 mM chloroacetamide (CAA, Sigma Aldrich Cat No: C0267) for 30 min at room temperature in the dark. Samples were diluted with 50 mM Tris.HCl, pH 8.5 to a final urea concentration of 2 M and treated with 1 µL PNGaseF (New England Biolabs,

Cat No P0705L) for 2 hrs at 35 ºC. Samples were then digested overnight at 35 ºC with 0.5 µg Trypsin/Lys-C (Mass Spectrometry grade, Promega Corporation, Cat No: V5072).

*Peptide Purification and Labeling*. Digestions were acidified with trifluoracetic acid (TFA, 0.5% v/v) and desalted on SPE columns (Pierce Cat no 89870) with a wash of 300 µL 0.1% TFA followed by elution in 70% acetonitrile 0.1% formic acid (FA).

*TMT labeling*. Peptides were dried by speed vacuum and resuspended in 24 µL of 50 mM triethylammonium bicarbonate pH 8.0 (TEAB) and labeled for two hours at room temperature with 0.2 mg of Tandem Mass Tag (TMT) reagent which was resuspended in 24 µL acetonitrile (Thermo Fisher Scientific, TMT™ Isobaric Label Reagent Set; Cat No 90111, lot no. WC306775 for 6-plex and WD320959 for 4-plex) [10]. Labelling reactions were quenched by adding 0.2% hydroxylamine (final v/v) to the reaction mixtures at room temperature for 15 minutes. Labeled peptides were then combined, mixed, and dried by speed vacuum. After drying, samples were resuspended in 0.1% TFA and desalted using Waters SepPak (Waters™ WAT054955) cartridges with a wash of 1 mL 0.1% TFA followed by elution in 70% acetonitrile 0.1% formic acid (FA). The specific TMT label corresponding to each WT and KO sample are designated in **Figure S9**.

*Phosphopeptide enrichment*. For the initial 6-plex of IP TMT samples, the entire sample was applied to a High-Select™ TiO2 Phosphopeptide enrichment tip (applied 3 times, washed and eluted as per manufacturer's instructions, Thermo Fisher Scientific, Cat No A32993). However, detected phosphopeptides from enriched and non-enriched fractions were combined in analysis due to limited phosphopeptide detection in the enriched fraction. Due to this, the phosphopeptide enrichment was not performed for the validatory 4-plex of IP TMT samples.

*Nano-LC-MS/MS Analysis*. Nano-LC-MS/MS analyses were performed on an EASY-nLC HPLC system (SCR: 014993, Thermo Fisher Scientific) coupled to Orbitrap Eclipse™ mass spectrometer (Thermo Fisher Scientific) with a FAIMS pro interface. Twenty and 25 % of each TMT mix was loaded onto a 25 cm aurora column (IonOpticks, AUR2-25075C18A) at 400 nL/min. 1/3 of the phosphopeptide fraction was injected. Peptides were eluted from 5-30% with mobile phase B (Mobile phases A: 0.1% FA, water; B: 0.1% FA, 80% Acetonitrile (Thermo Fisher Scientific Cat No: LS122500)) over 160 minutes, 30-80% B over 10 mins; and dropping from 80-10% B over the final 10 min. The mass spectrometer was operated in positive ion mode with 3 FAIMS CVs (-45, -55, -70) and 1.3 sec cycle time per CV. Data-dependent acquisition method with advanced peak determination and Easy-IC (internal calibrant) were used. Precursor scans (m/z 400-1600) were done with an orbitrap resolution of 120000, RF lens% 30, maximum inject time 105 ms, standard AGC target, MS2 intensity threshold of 2.5e4, including charges of 2 to 6 for fragmentation with 60 sec dynamic exclusion. MS2 scans were performed with a quadrupole isolation window of 0.7 m/z, 38% HCD CE, 50000 resolution, 200% normalized AGC target, dynamic maximum IT fixed first mass of 100 m/z. The data were recorded using Thermo Fisher Scientific Xcalibur (4.3) software (Thermo Fisher Scientific Inc.).

*Mass spectrometry data analysis*. Resulting RAW files were analyzed in Proteome Discover™ 2.5 (Thermo Fisher Scientific, RRID: SCR_014477) with a *Mus musculus* UniProt FASTA (both reviewed and unreviewed sequences) plus common contaminants (49922 total sequences). Quantification methods utilized isotopic impurity levels available from Thermo Fisher Scientific. SEQUEST HT searches were conducted with a maximum number of 3 missed cleavages; precursor mass tolerance of 10 ppm; and a fragment mass tolerance of 0.02 Da. Static modifications used for the search were carbamidomethylation on cysteine (C) residues. Dynamic modifications used for the search were oxidation of methionines, TMT label on the N-termini of peptides, TMT label on lysine (K) residues, phosphorylation (S, T, Y) and deamidation of N (max 3 dynamic mods). Dynamic Protein terminus modifications allowed were: acetylation (N-terminus), Met-loss or Met-loss plus acetylation (N-terminus). Percolator False Discovery Rate was set to a strict setting of 0.01 and a relaxed setting of 0.05. IMP-ptm-RS node was used for all modification site localization scores. In the consensus workflows, data were normalized using a fasta file of all 3 GRM5 isoforms. Co-isolation thresholds of 50% and average reporter ion S/N cutoffs of 10 were used for quantification. Lot specific isotopic impurity levels were corrected for. Resulting normalized abundance values for each sample type, abundance ratio and log2(abundance ratio) values; and respective p-values (t-test) from Proteome Discover™ were exported to Microsoft Excel. All processed and raw data are uploaded to ProteomeXchange (accession number pending) [11, 12].

<u>Statistics</u>

Rotarod performance was analyzed by two-way analysis of variance (ANOVA) with repeated measures with post-hoc Dunnett's multiple comparisons test and paired t-tests. Amphetamine-induced locomotor sensitization was analyzed by three-way ANOVAs with repeated measures. SAPAP3 WT and KO behavior in OF was analyzed by two-way ANOVAs with post-hoc Šídák's or Tukey's multiple comparisons test and Pearson's correlation analyses. mGluR5 PAM behavior in OF was analyzed by two-way ANOVAs with post-hoc Šídák's multiple comparisons test, Pearson's correlation, unpaired t-tests, and one-way ANOVAs with post-hoc Dunnett's multiple comparisons test. Field electrophysiology recordings were analyzed by two-way ANOVA with repeated measures, one-way ANOVA, or one-way ANOVAs with repeated measures with post-hoc Dunnett's multiple comparisons tests. Immunoblots were quantified as previously described [2] and analyzed by unpaired t-test or one-way ANOVAs with post-hoc Dunnett's multiple comparisons test. Quantitative proteomics results were first analyzed by $log_2$ fold-change (KO/WT), then one-tailed unpaired t-tests were performed. String-db was used to generate protein-protein interaction networks and perform gene ontology (GO) analyses, which used false discover rate to filter significant GO terms. In all ANOVA tests described above, post-hoc tests were only performed when column, row, or interaction p-value $< 0.05$. If $p < 0.05$, multiple comparisons were only performed within columns and/or rows that had a significant ANOVA effect. For t-tests of proteomics phosphorylation and interaction data, $p < 0.1$ was used.

### References

1. Edler, M.C., et al., *Mechanisms Regulating the Association of Protein Phosphatase 1 with Spinophilin and Neurabin.* ACS Chem Neurosci, 2018. **9**(11): p. 2701-2712.
2. Morris, C.W., et al., *The association of spinophilin with disks large-associated protein 3 (SAPAP3) is regulated by metabotropic glutamate receptor (mGluR) 5.* Mol Cell Neurosci, 2018. **90**: p. 60-69.
3. Kalueff, A.V., et al., *Analyzing grooming microstructure in neurobehavioral experiments.* Nat Protoc, 2007. **2**(10): p. 2538-44.
4. Kalueff, A.V., et al., *Neurobiology of rodent self-grooming and its value for translational neuroscience.* Nat Rev Neurosci, 2016. **17**(1): p. 45-59.
5. Yin, H.H., et al., *Dynamic reorganization of striatal circuits during the acquisition and consolidation of a skill.* Nat Neurosci, 2009. **12**(3): p. 333-41.
6. Salek, A.B., et al., *Spinophilin regulates phosphorylation and interactions of the GluN2B subunit of the N-methyl-d-aspartate receptor.* J Neurochem, 2019. **151**(2): p. 185-203.
7. Hiday, A.C., et al., *Mechanisms and Consequences of Dopamine Depletion-Induced Attenuation of the Spinophilin/Neurofilament Medium Interaction.* Neural Plast, 2017. **2017**: p. 4153076.
8. Watkins, D.S., et al., *Proteomic Analysis of the Spinophilin Interactome in Rodent Striatum Following Psychostimulant Sensitization.* Proteomes, 2018. **6**(4).
9. Grecco, G.G., et al., *A multi-omic analysis of the dorsal striatum in an animal model of divergent genetic risk for alcohol use disorder.* J Neurochem, 2021. **157**(4): p. 1013-1031.
10. Li, J., et al., *TMTpro reagents: a set of isobaric labeling mass tags enables simultaneous proteome-wide measurements across 16 samples.* Nat Methods, 2020. **17**(4): p. 399-404.
11. Kelleher, R.J., 3rd, et al., *High-throughput sequencing of mGluR signaling pathway genes reveals enrichment of rare variants in autism.* PLoS One, 2012. **7**(4): p. e35003.
12. Deutsch, E.W., et al., *The ProteomeXchange consortium in 2020: enabling 'big data' approaches in proteomics.* Nucleic Acids Res, 2020. **48**(D1): p. D1145-d1152.

### Figure S1

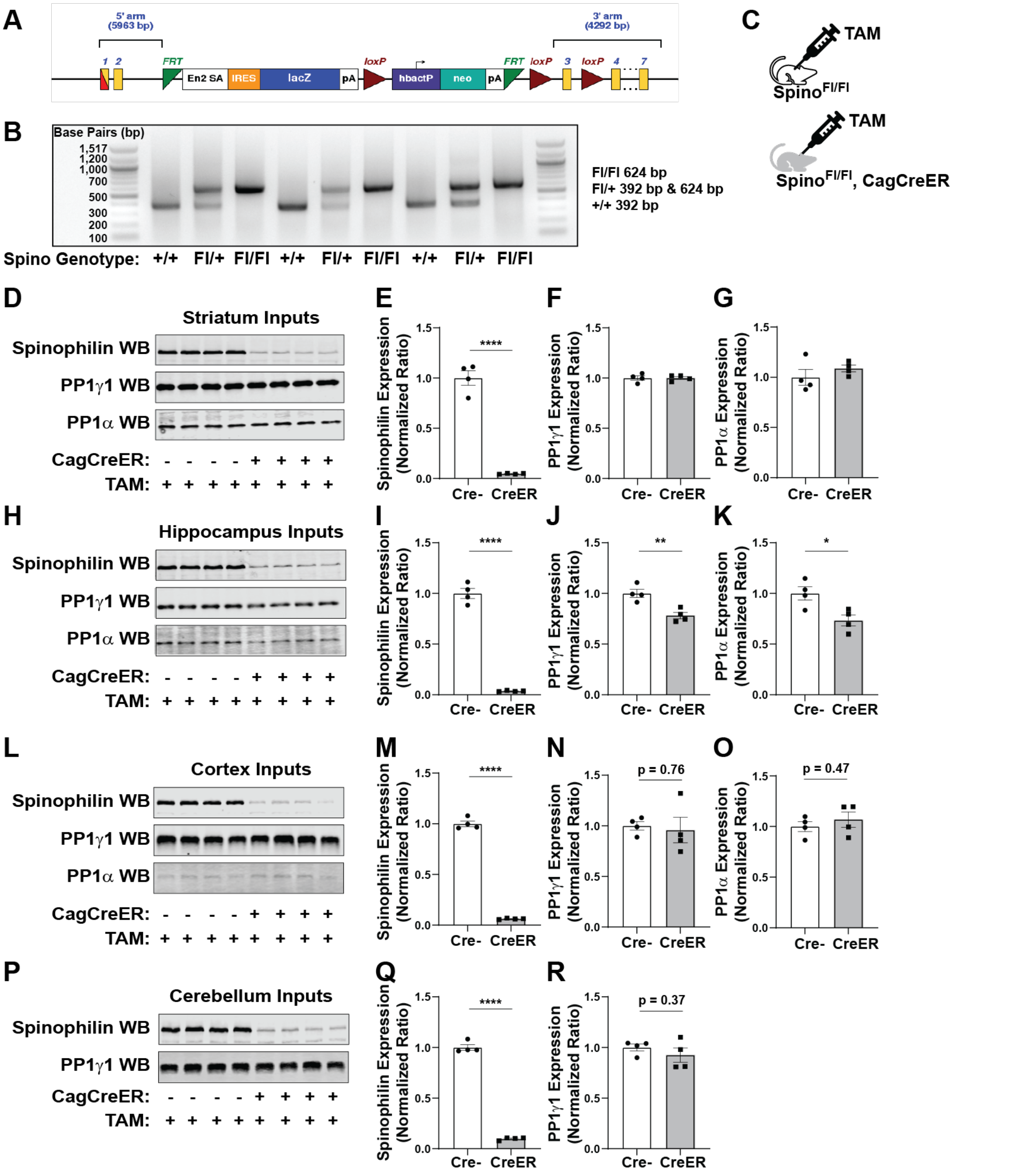


**Figure S1: Biochemical validation of conditional spinophilin knockout mice.** **A)** Design of targeted DNA insert used to create spinophilin gene conditional line. **B)** Representative example of genotyping +/+, Fl/+, and Fl/Fl spinophilin alleles. **C)** Conditional spinophilin KO mice were crossed with an inducible Cre recombinase (Cre) line expressed under the Cag promotor (CagCreER). Male and female Spino$^{Fl/Fl}$ mice

+/- CagCreER were given 5 daily I.P. injections of tamoxifen (TAM, 24mg/kg) and whole striatum was dissected 28 days later for semi-quantitative immunoblot analysis of **D-G)** striatal, **H-K)** hippocampus, **L-O)** cortex, and **P-R)** cerebellum inputs. Student's t-tests were performed to determine the effect CagCreER has on protein expression. CagCreER significantly decreased **E)** spinophilin ($p<0.0001$) but not **F)** PP1$\gamma$1 ($p=0.98$) or **G)** PP1$\alpha$ ($p=0.34$) protein expression in striatum. In hippocampus, CagCreER significantly decreased **I)** spinophilin ($p<0.0001$), **J)** PP1$\gamma$1 ($p=0.005$), and **K)** PP1$\alpha$ ($p=0.02$) expression, suggesting there may be compensatory changes in PP1 levels similar to Spino$^{-/-}$ hippocampus [6]. In cortex, CagCreER significant decreased **M)** spinophilin ($p<0.0001$) but not **N)** PP1$\gamma$1 ($p=0.76$) or **O)** PP1$\alpha$ ($p=0.47$) expression. In cerebellum, CagCreER significantly decreased **Q)** spinophilin ($p<0.0001$) but not **K)** PP1$\gamma$1 ($p=0.37$). Consistent with in-situ hybridization data reported in the Allen Brain Atlas, we detected minimal PP1$\alpha$ expression in cerebellum (data not shown). N=4 TAM-treated Spino$^{Fl/Fl}$ (2 male), 4 TAM-treated Spino$^{Fl/Fl}$/CagCreER (2 male). Data ± SEM. *$p\leq0.05$, **$p\leq0.01$, ****$p<0.0001$.

### Figure S2

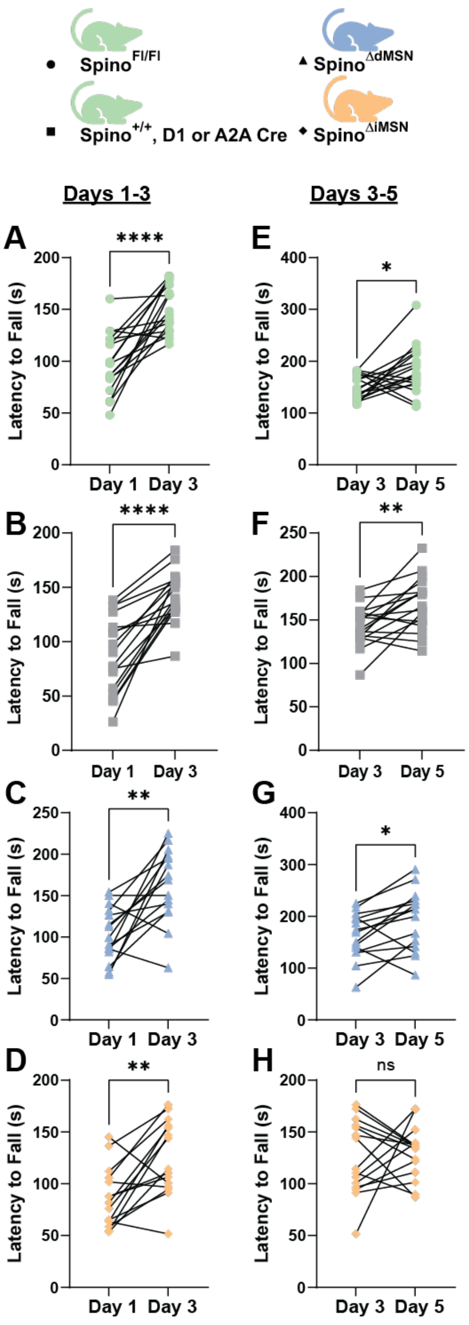


**Figure S2: Early- and late-stage performance on accelerating rotarod.** Rotarod performance for Spino$^{Fl/Fl\ or\ Fl/+}$, Spino$^{+/+}$-D1 Cre or -A2A Cre, Spino$^{\Delta dMSN}$, and Spino$^{\Delta iMSN}$ mice was broken down into early stage (days 1-3) and late-stage (days 3-5) as described in Yin et al, 2009 [5]. All genotypes increased rotarod performance from day 1 to 3 **(A-D)**, however, unlike controls and spino$^{\Delta dMSN}$, rotarod performance plateaued from days 3-5 (**E-H)** in Spino$^{\Delta iMSN}$ mice. N=17 Spino$^{Fl/Fl}$ (7 male), 17 Spino$^{+/+}$-D1 or -A2A Cre (11 male), 15 Spino$^{\Delta dMSN}$ (10 male), 15 Spino$^{\Delta iMSN}$ (5 male).

**Figure S3**

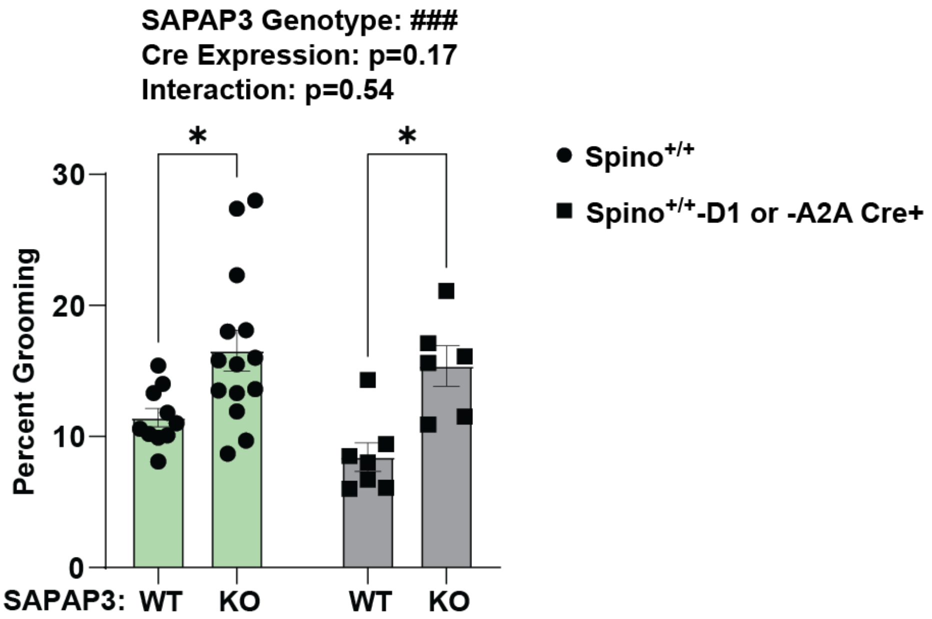


**Figure S3: D1- and A2A-Cre expression do not decrease excessive grooming in SAPAP3 knockout mice.** Percent grooming behavior of SAPAP3 WT and KO mice expressing wild type spinophilin (Spino$^{+/+}$) and D1- or A2A-Cre and SAPAP3 WT and KO mice Cre- mice was measured for 1-hour at 8-weeks of age. Two-way ANOVA with a post-hoc Šídák's multiple comparisons test detected a significant SAPAP3 genotype effect in Cre- (p=0.014) and D1- or A2A Cre+ (p=0.013) mice. N=10 SAPAP3 WT (6 male), 14 SAPAP3 KO (5 male), 7 SAPAP3 WT/D1- or A2A-Cre (4 male), 6 SAPAP3 KO/D1- and A2A-Cre (4 male). Data ± SEM. Significant two-ANOVA effects denoted by ###p≤0.001. Significant post-hoc tests denoted by *p≤0.05.

**Figure S4**

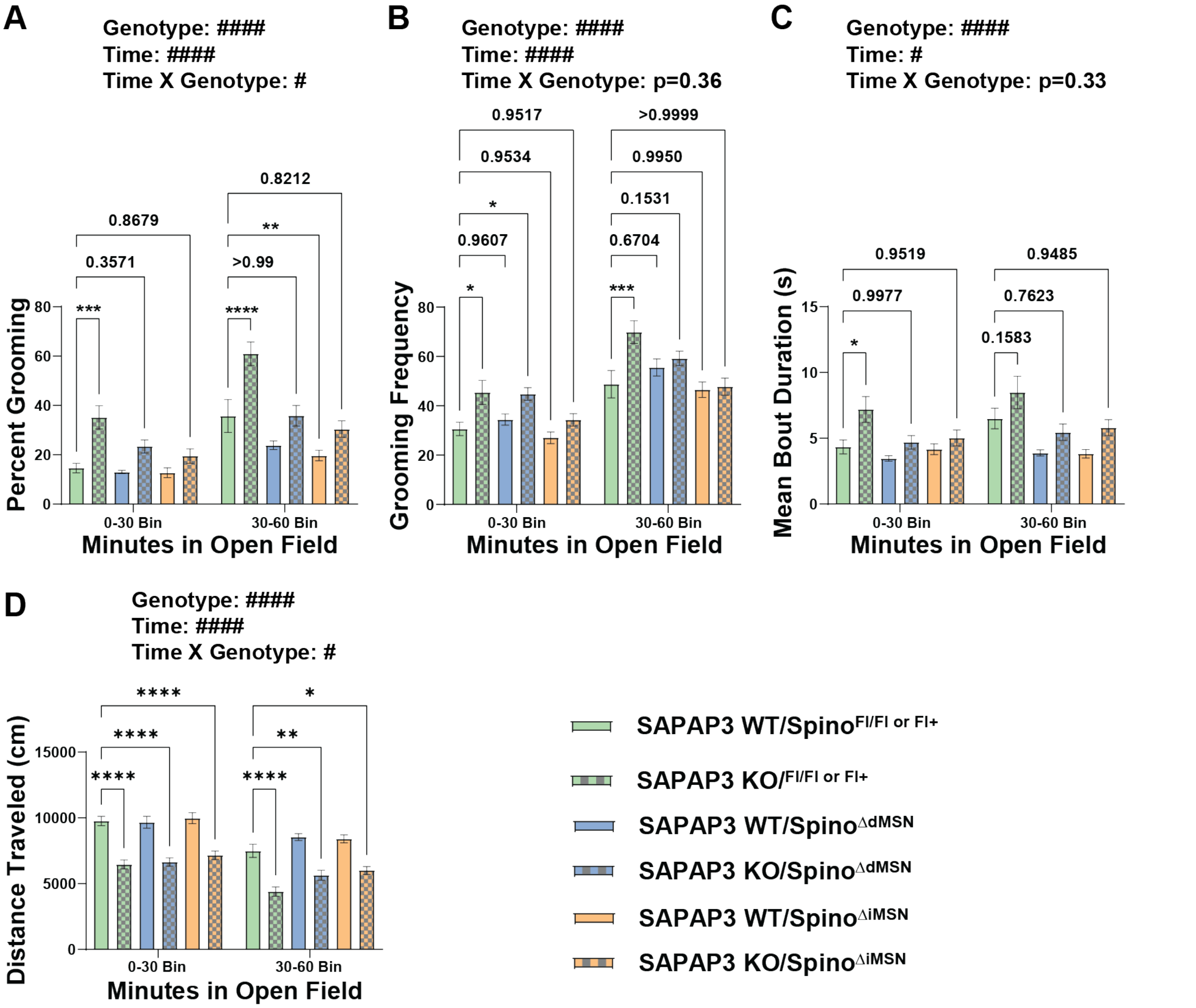


**Figure S4: MSN-specific spinophilin knockout effects on SAPAP3 knockout motor dysfunction broken into 30-minute bins.** The 60-minute open field behavior measurement of SAPAP3 WT and KO/MSN-specific spinophilin knockout was halved into 30-minute bins. Two-way ANOVAs with repeated measures were performed to determine the effect time and genotype have on **A)** percent grooming, **B)** grooming frequency, **C)** mean grooming bout duration, and **D)** distance traveled. N=11-13 SAPAP3 WT-KO/Spino$^{Fl/Fl\text{ or }Fl/+}$ (7-5 male), N=9-12 SAPAP3 WT-KO/Spino$^{\Delta dMSN}$ (4-6 male), and N=12-12 SAPAP3 WT-KO/Spino$^{\Delta iMSN}$ (4-6 male). Data ± SEM. Significant two-ANOVA effects denoted by #p≤0.05 and ####p<0.0001. Significant post-hoc tests denoted by *p≤0.05, **p≤0.01, ***p≤0.001, ****p<0.0001.

**Figure S5**

**A**

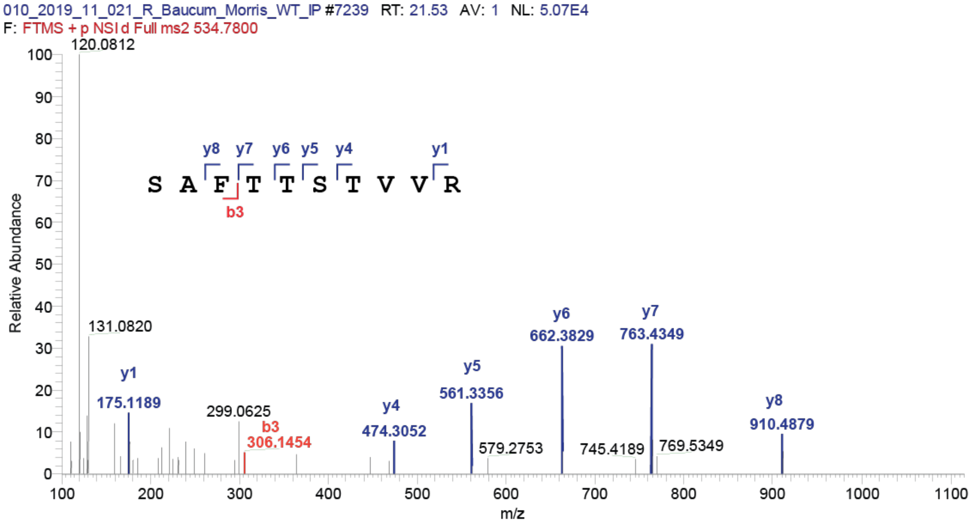


**B**

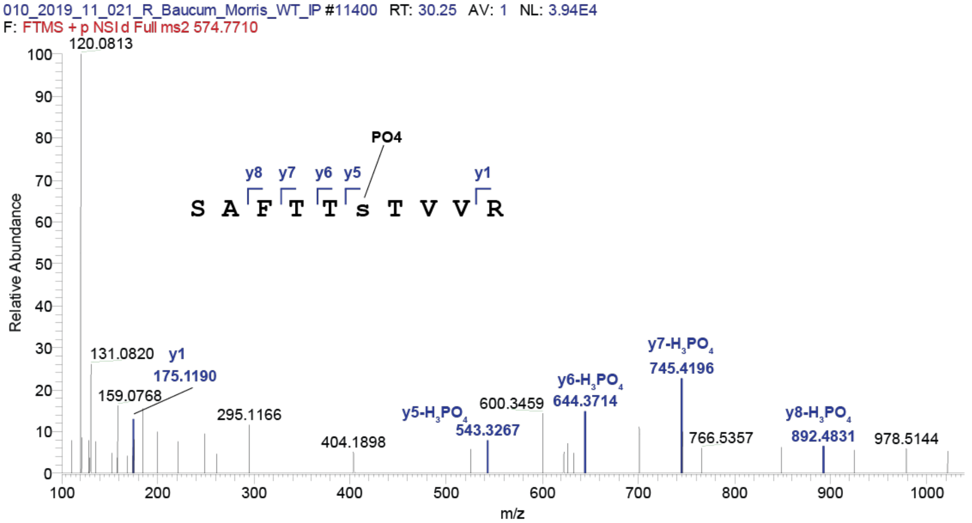


**Figure S5: Mass spectrum of mGluR5 Ser839 phosphopeptide.** MS/MS spectrum hand annotated with b- and y-series ions of **A)** total- and **B)** phospho-mGluR5 peptide containing Ser839.

**Figure S6**

**A**

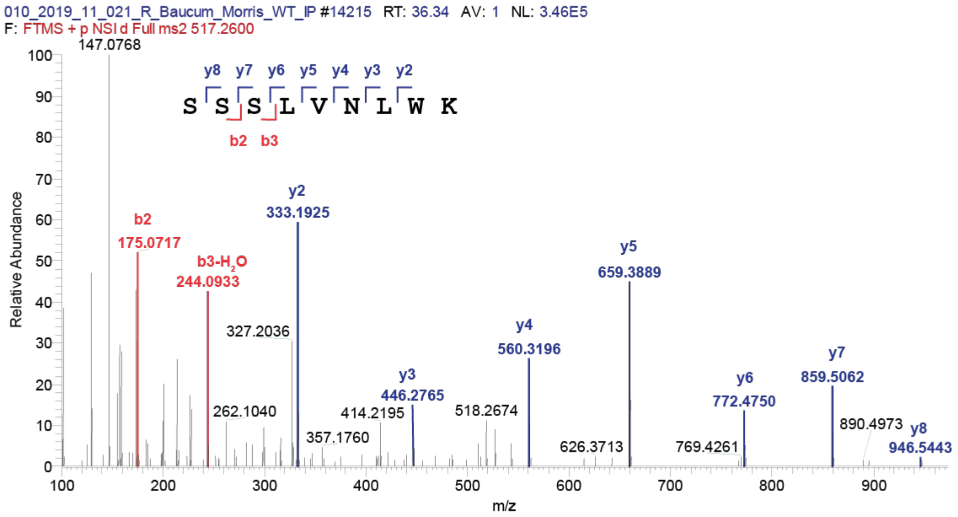


**B**

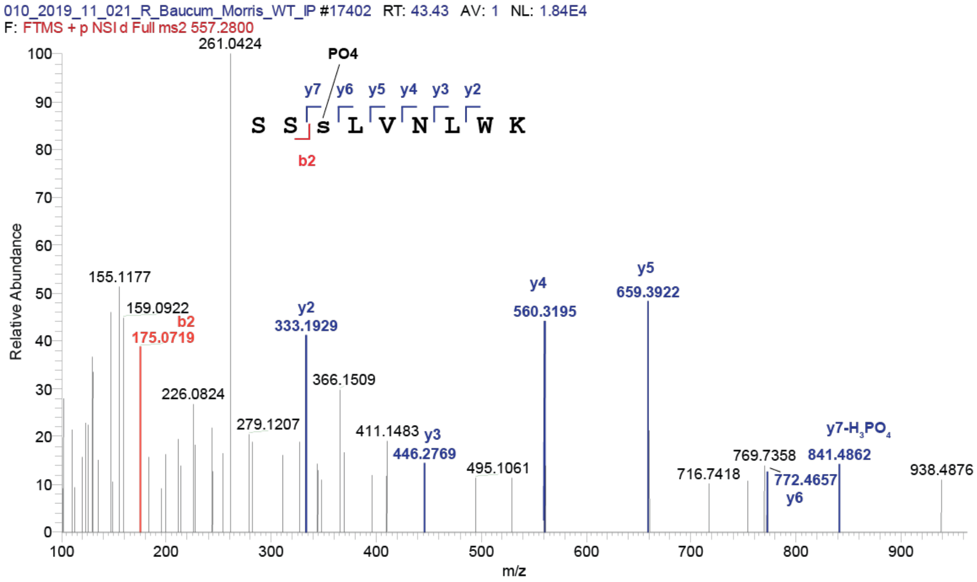


**Figure S6: Mass spectrum of mGluR5 Ser860 phosphopeptide.** MS/MS spectrum hand annotated with b- and y-series ions of **A)** total- and **B)** phospho-mGluR5 peptide containing Ser860.

**Figure S7**

**A**

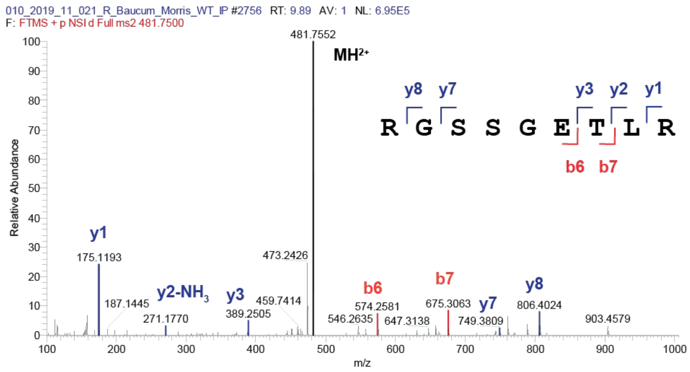


**B**

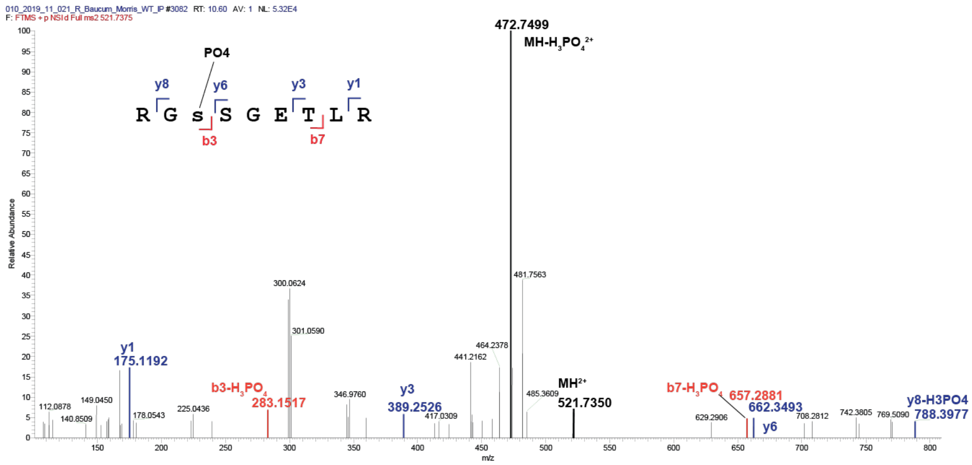


**Figure S7: Mass spectrum of mGluR5 Ser870 phosphopeptide.** MS/MS spectrum hand annotated with b- and y-series ions of **A)** total- and **B)** phospho-mGluR5 peptide containing Ser870.

**Figure S8**

**A**

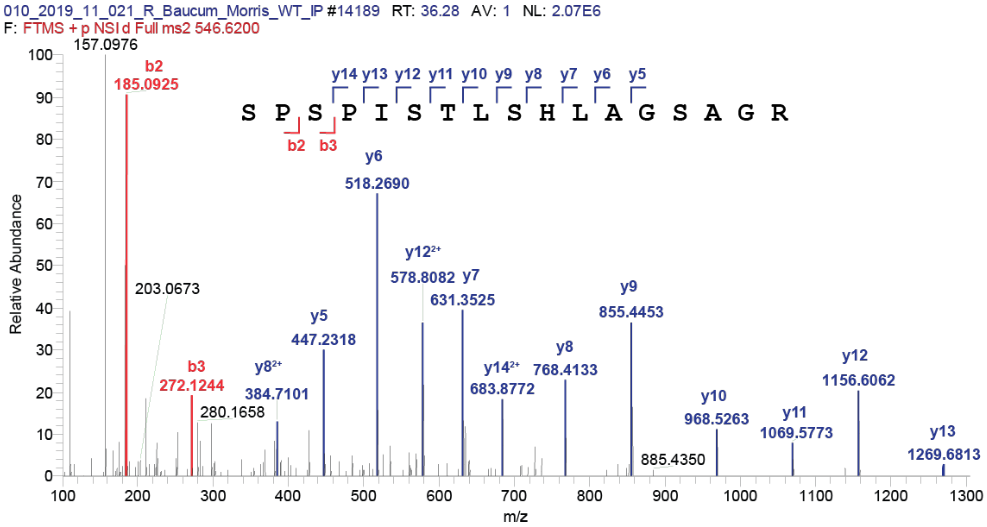


**B**

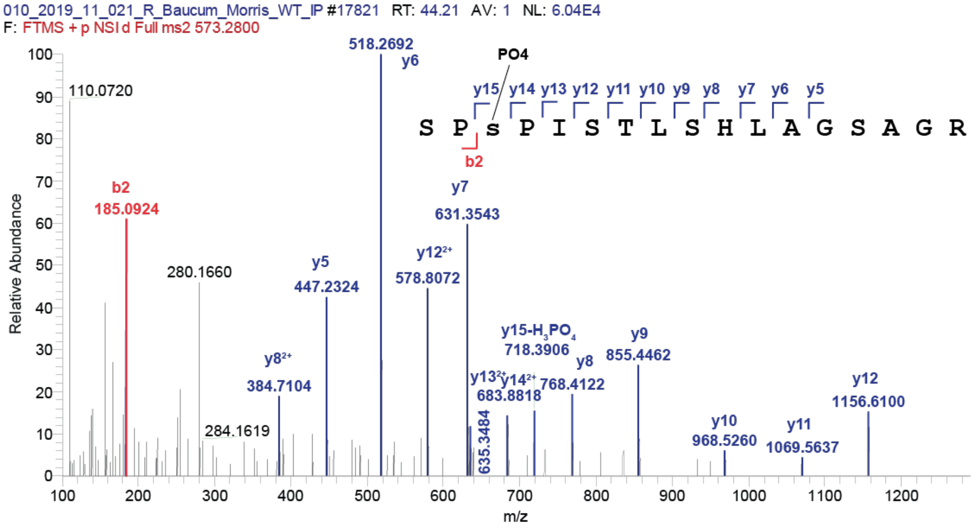


**Figure S8: Mass spectrum of mGluR5 Ser1016 phosphopeptide.** MS/MS spectrum hand annotated with b- and y-series ions of **A)** total- and **B)** phospho-mGluR5 peptide containing Ser1016.

**Figure S9**

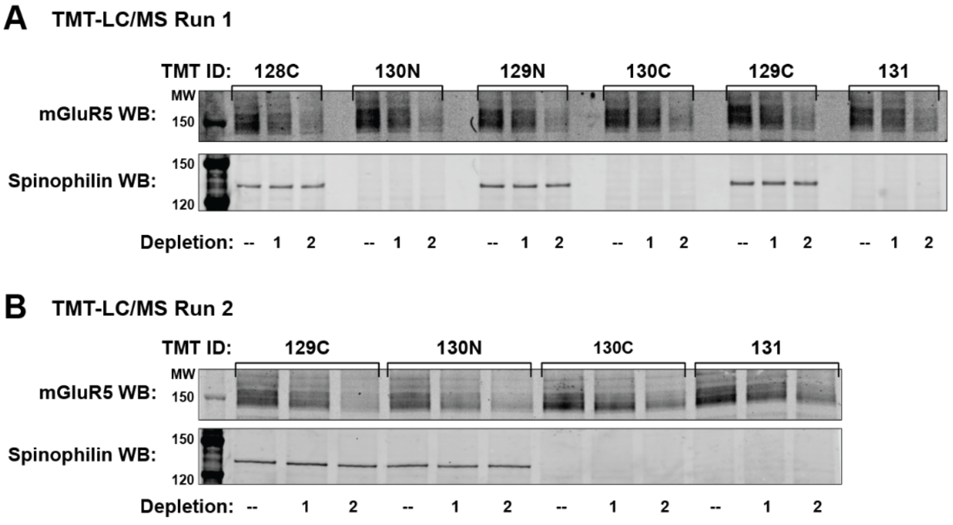


**Figure S9: Immunoblot validation of samples submitted for TMT-LC/MS run 1 and 2.** Western blots confirming sequential mGluR5 immunodepletion and spinophilin expression from TMT-LC/MS **A)** run1 and **B)** run 2. TMT IDs above lanes corresponds to isobaric tags listed in Table S2 (run 1) and Table S5 (run 2).

**Figure S10**

**Log₂(Fold-Change (KO/WT) < -0.2**

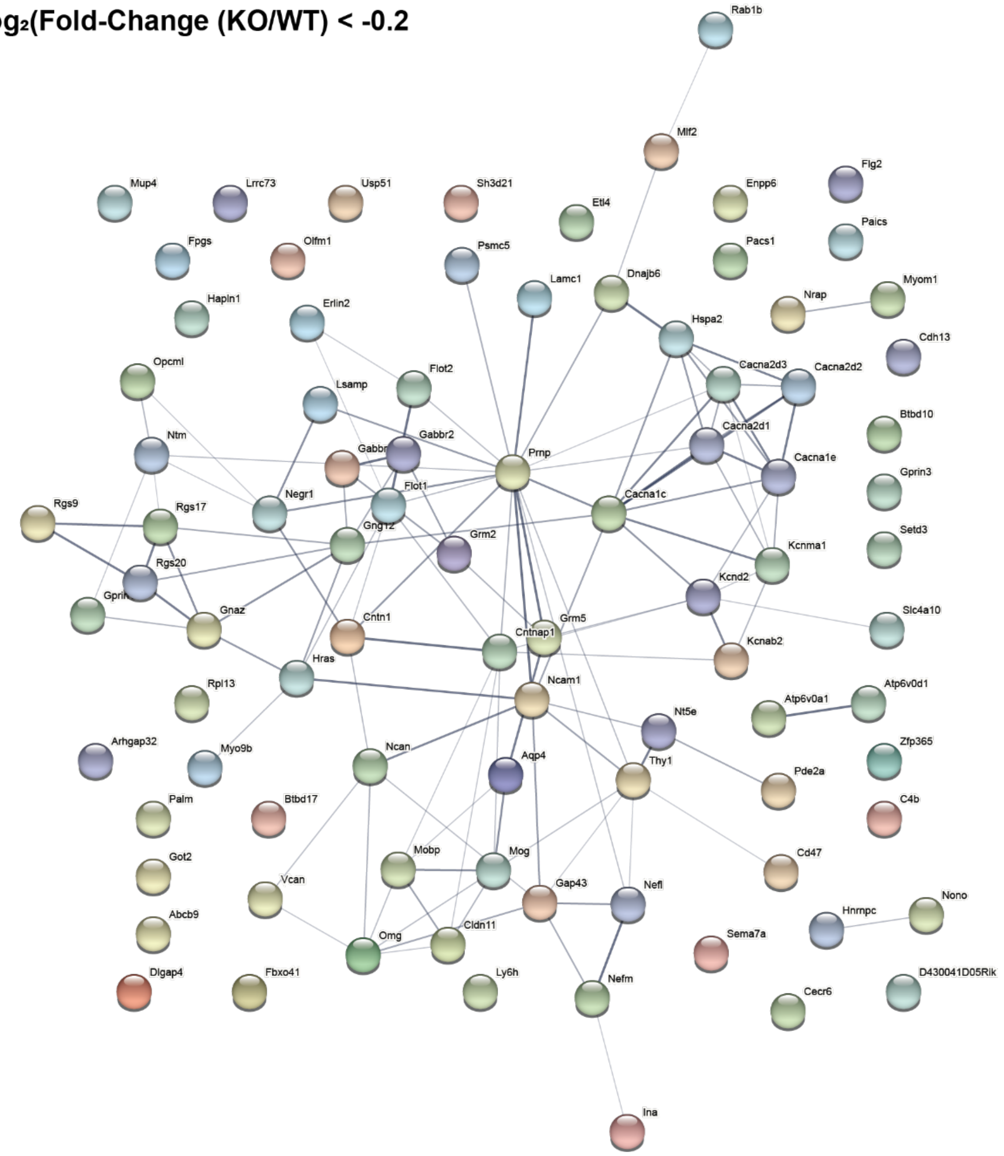


**Figure S10: Complete PPI network of decreased mGluR5 interactions.** All 92 decreased mGluR5 interactors ($log_2$-fold change < -0.2) were graphed in string-DB to visualize how loss of spinophilin modulates mGluR5's protein-protein interaction (PPIs) network.

### Figure S11

**$Log_2$(Fold-Change (KO/WT) > 0.2**

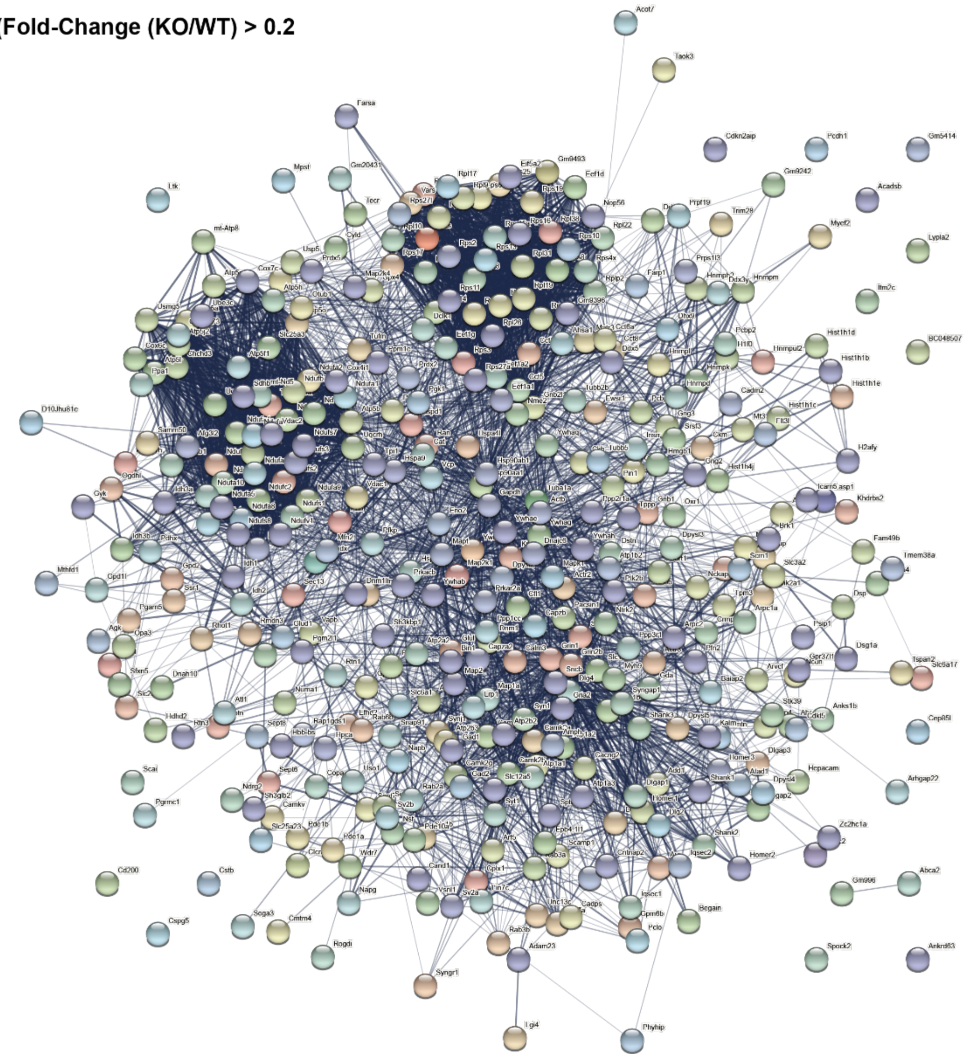

**Figure S11: Complete PPI network of increased mGluR5 interactors.** All 426 increased mGluR5 interactors ($log_2$-fold change > 0.2) were graphed in string-DB to visualize how loss of spinophilin modulates mGluR5's protein-protein interaction (PPIs) network.

**Figure S12**

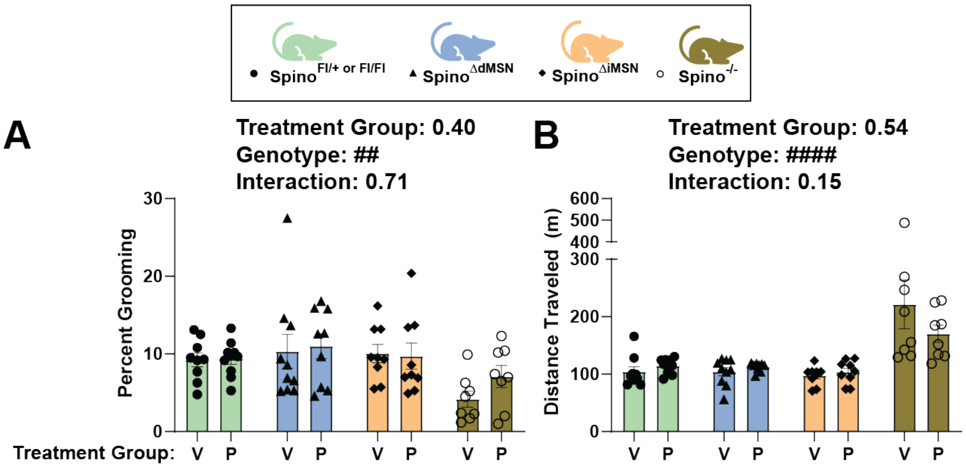


**Figure S12: Pre-injection open field behavior prior to treatment with mGluR5 PAM VU'172 (20 mg/kg).** Two-way ANOVA with post-hoc Dunnett's multiple comparisons test determined there were no pre-existing treatment group effects on **A)** percent grooming or **B)** distance traveled in the 30-minutes pre-injection period (Figure 5A), however, percent grooming was significantly decreased ($p=0.04$), and distance traveled was significantly increased ($p<0.0001$) in spino$^{-/-}$ compared to control. N=9V/10P Spino$^{Fl/+ \text{ or } Fl/Fl}$ (3/3 male), 10V/9P Spino$^{\Delta dMSN}$ (5/5 male), and 9V/9P Spino$^{\Delta iMSN}$ (4/4 male), 8V/8P Spino$^{-/-}$ (3/4 male). Significant two-way ANOVA effects denoted by ##$p\leq0.01$ and ####$p<0.0001$.

**Figure S13**

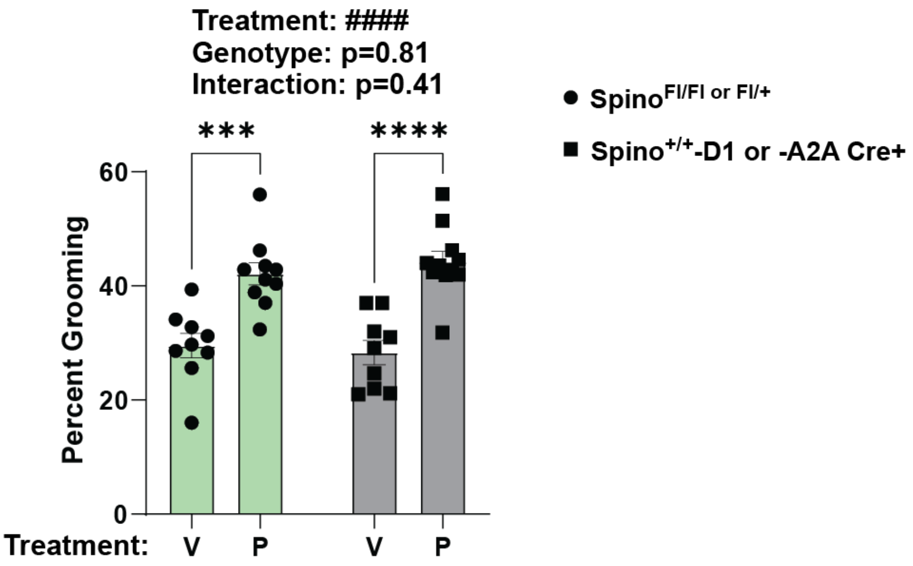


**Figure S13: D1- or A2A-Cre expression don't affect VU'172 grooming.** Percent grooming from vehicle- and VU'172-treated Spino$^{Fl/Fl\ or\ Fl/+}$ control group (Figure 5A) was compared to vehicle- and VU'172-treated Spino$^{+/+}$-D1 or -A2A Cre control group. Two-way ANOVA with post-hoc Šídák's multiple comparisons test determined a significant VU'172 treatment effect in Spino$^{Fl/Fl\ or\ Fl/+}$ control and Spino$^{+/+}$-D1 or -A2A Cre control groups ($p=0.0002$ and $p<0.0001$, respectively). N=9V/10P Spino$^{Fl/+\ or\ Fl/Fl}$ (3/3 male) and 9V/11P Spino$^{+/+}$-D1 Cre or -A2A Cre (3/4 male). Data ± SEM. Significant two-ANOVA effects denoted by ####$p<0.0001$. Significant post-hoc tests denoted by ***$p\leq0.001$ and ****$p<0.0001$.

**Figure S14**

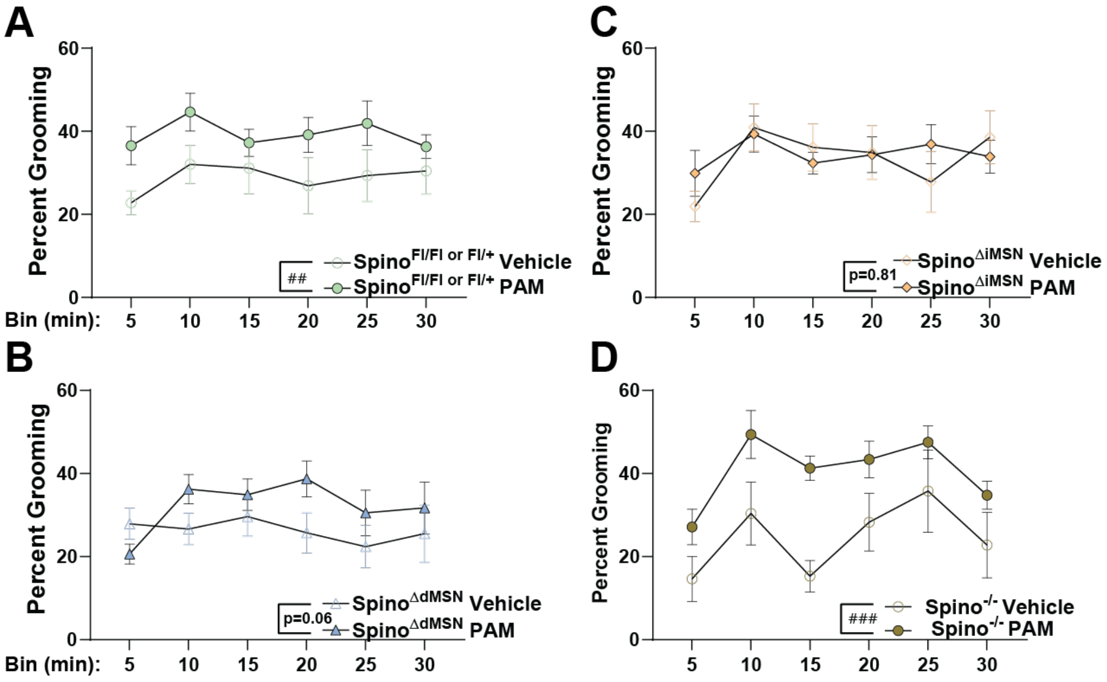


**Figure S14: Post-injection percent grooming following vehicle or mGluR5 PAM (20 mg/kg) treatment in 5-minute bins by genotype.** Percent grooming response from vehicle- or VU'172-treated **A)** Spino$^{Fl/Fl\ or\ Fl/+}$, **B)** Spino$^{\Delta dMSN}$, **C)** Spino$^{\Delta iMSN}$ , and Spino$^{-/-}$ was broken into 5-minute bins. Two-way ANOVAs with repeated measures analyses determined significant VU'172 treatment effect on percent grooming in Spino$^{Fl/Fl\ or\ Fl/+}$ control (p=0.001) and Spino$^{-/-}$ (p=0.0003), but not Spino$^{\Delta dMSN}$ (p=0.06) or Spino$^{\Delta iMSN}$ (p=0.81) groups. N=9V/10P Spino$^{Fl/+\ or\ Fl/Fl}$ (3/3 male), 10V/10P Spino$^{\Delta dMSN}$ (5/6 male), and 9V/9P Spino$^{\Delta iMSN}$ (4/4 male), 8V/8P Spino$^{-/-}$ (3/4 male). Data ± SEM. Significant two-ANOVA effects denoted by ##p≤0.01 and ###p≤0.001.

### Figure S15

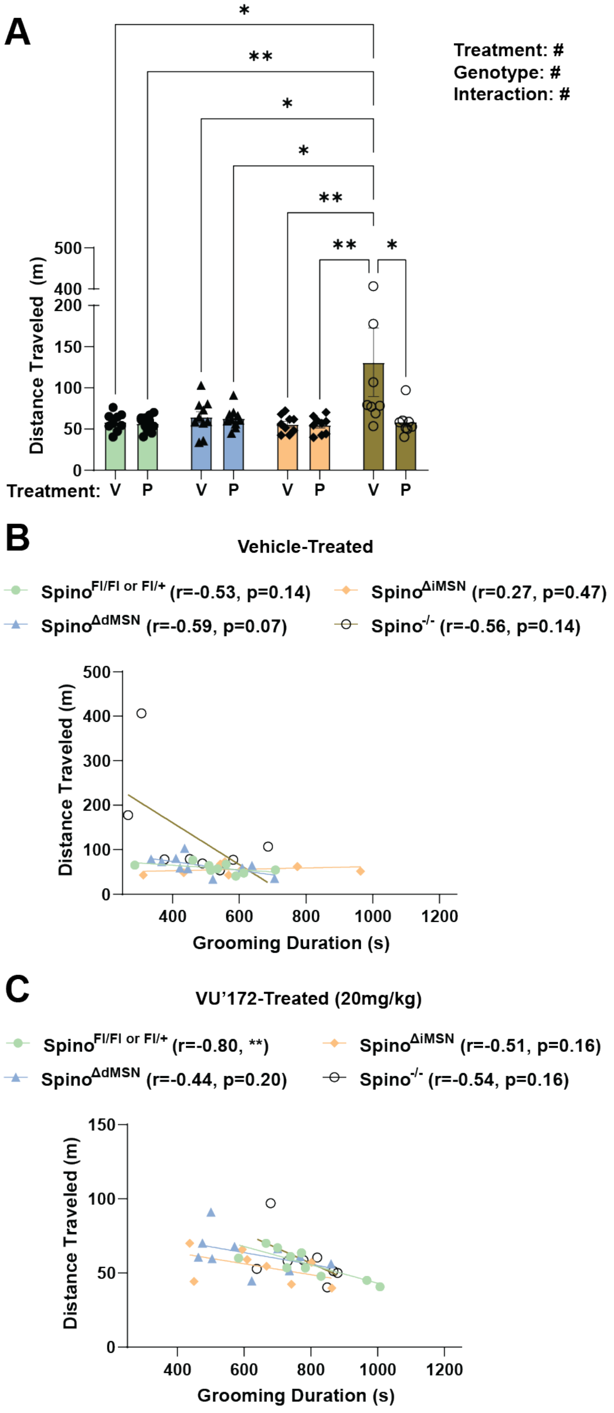


**Figure S15: VU'172 effects on locomotion in OF. A)** Two-way ANOVA with post-hoc Dunnett's multiple comparisons test determined distance traveled in vehicle-treated spino$^{-/-}$ mice was significantly increased compared to all other groups. Person's correlation analysis did not detect a significant correlation between grooming duration and distance traveled in **B)** vehicle-treated groups, and a significant negative correlation was only detected in **C)** VU'172-treated Spino$^{Fl/Fl\ or\ Fl/+}$ control mice. N=9V/10P Spino$^{Fl/+\ or\ Fl/Fl}$

(3/3 male), 10V/10P Spino$^{\Delta dMSN}$ (5/6 male), and 9V/9P Spino$^{\Delta iMSN}$ (4/4 male), 8V/8P Spino$^{-/-}$ (3/4 male). Data ± SEM. Significant two-ANOVA effects denoted by #p≤0.05. Significant post-hoc tests denoted by *p≤0.05, **p≤0.01.

**Figure S16**

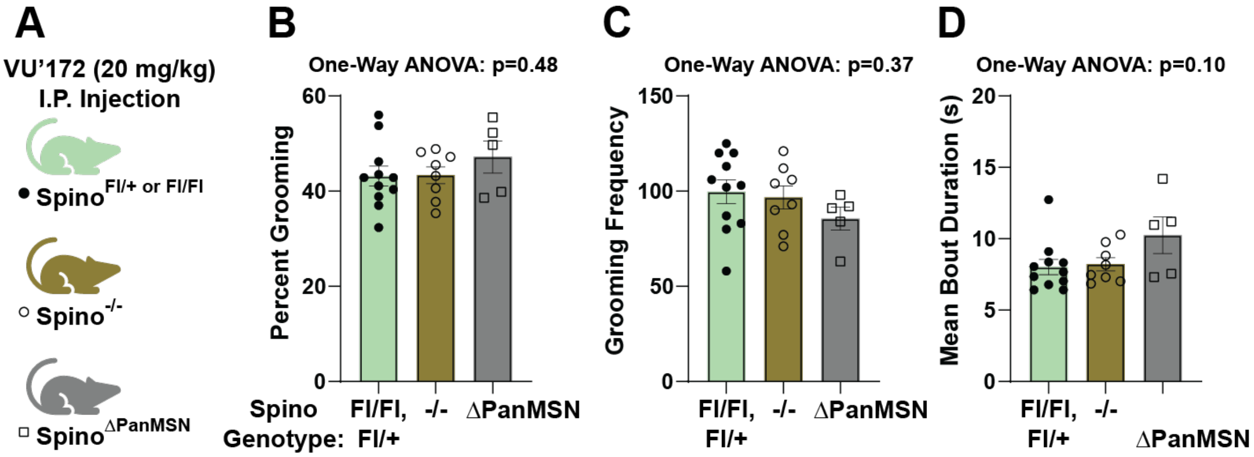


**Figure S16: Depletion of spinophilin from dMSNs and iMSNs, together, doesn't decrease VU'172 grooming behavior. A)** Spino$^{\Delta PanMSN}$ were treated with mGluR5 PAM (VU'172, P) (20 mg/kg) and compared with existing Spino$^{Fl/Fl\ or\ Fl/+}$ control and Spino$^{-/-}$ data (Figure 5B). One-way ANOVA tests determined spino$^{\Delta PanMSN}$ doesn't impact **B)** percent grooming (p=0.48), **C)** grooming frequency (p=0.37), nor **D)** mean grooming bout duration (p=0.10). N=10P Spino$^{Fl/+\ or\ Fl/Fl}$ (3 male), 8P Spino$^{-/-}$ (3/4 male), and 5P Spino$^{\Delta PanMSN}$ (4 male). Data ± SEM.

**Figure S17**

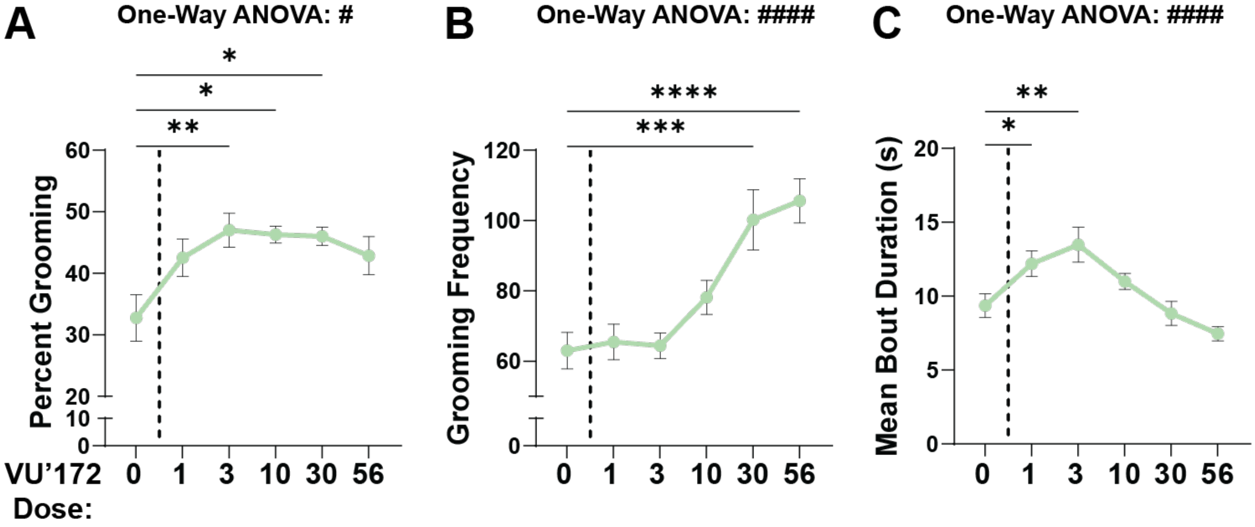


**Figure S17: VU'172 dose response in Spino$^{Fl/Fl}$ control mice.** Spino$^{Fl/Fl}$ controls were treated with vehicle, 1, 3, 10, 30, or 56 mg/kg VU'172 and grooming behavior was measured as described in Figure 5A. One-way ANOVAs with post-hoc Dunnett's multiple comparison's test determined that **A)** percent grooming is significantly increased at 3 mg/kg (p=0.003), 10 mg/kg (p=0.011), and 30 mg/kg (p=0.017), whereas 1 mg/kg (p=0.06) and 56 mg/kg (p=0.056) VU'172 doses were trending toward a significant increase. **B)** Grooming frequency had a classical dose-response curve that significantly increased from vehicle control at 30 mg/kg (p=0.003) and 56 mg/kg (p<0.0001) VU'172. Alternatively, **C)** Mean grooming bout duration, had an inverted U-shaped dose-response curves that is significantly increased at 1 mg/kg (p=0.040) and 3 mg/kg (p=0.005) VU'172, but not 10 mg/kg, 30 mg/kg, or 56 mg/kg (p=0.52, p=0.98, p=0.28, respectively). N=8-13 Spino$^{Fl/Fl}$ (6-9 male). Data ± SEM. Significant one-ANOVA effects denoted by #p≤0.05 and ####p<0.0001. Significant post-hoc tests denoted by *p≤0.05, **p≤0.01, ****p<0.0001.

### Figure S18

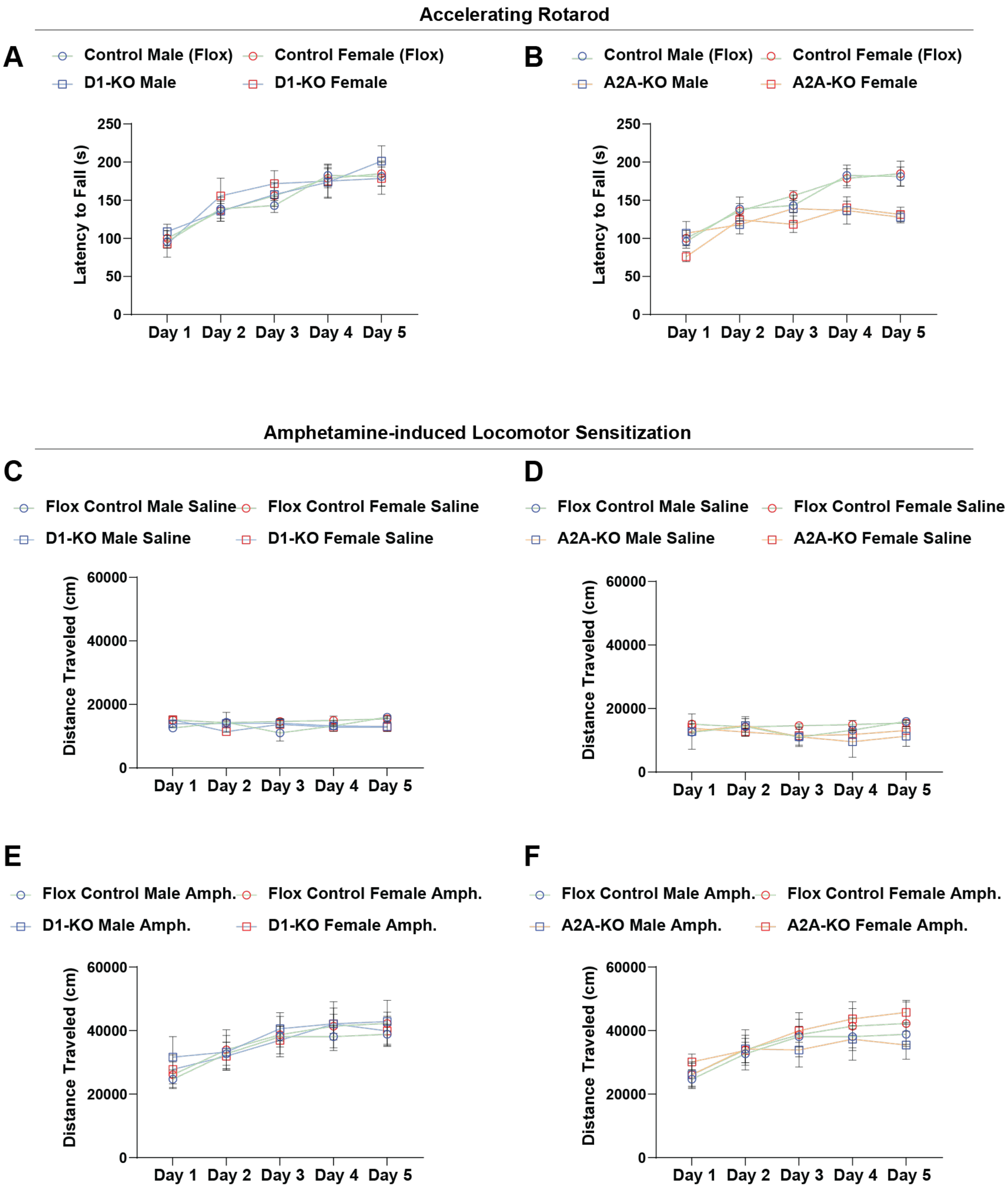


**Figure S18: Rotarod and amphetamine-induced locomotor sensitization by sex.** Male (blue points) and female (red points) control ($Spino^{Fl/Fl}$ and $Spino^{+/+}$-D1 or -A2A Cre) (green bars), $Spino^{\Delta dMSN}$ (blue bars), and $Spino^{\Delta iMSN}$ (orange bars) were plotted separately for **A-B.** latency to fall, **C-D.** saline-treated distance traveled, and **E-F.** amphetamine-treated distance traveled. N=5-11 for rotarod. Data ± SEM. N=2-10.

**Figure S19**

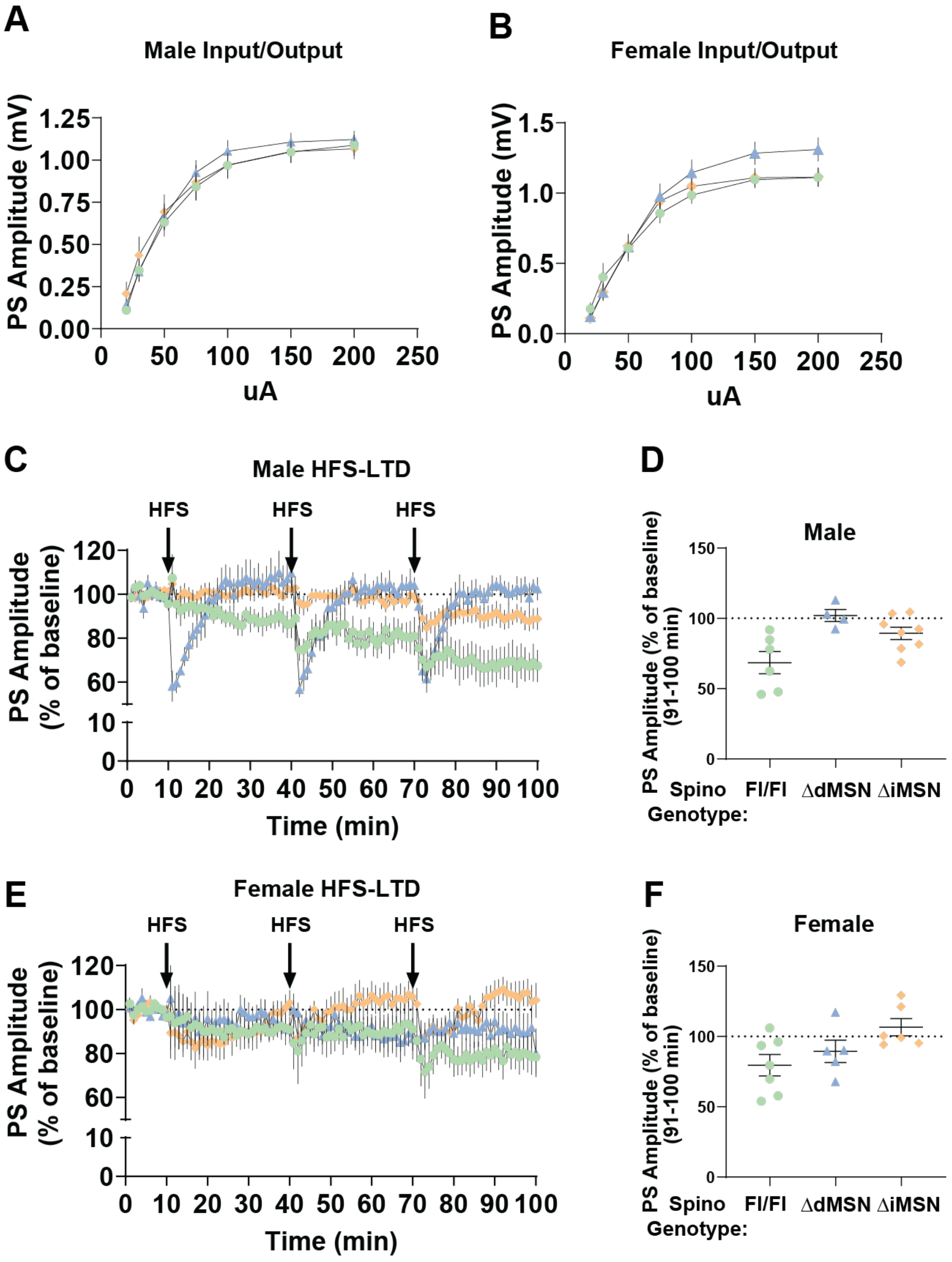


**Figure S19: Dorsolateral striatum field electrophysiology recordings by sex. A)** Male and **B)** female input/output field population spike amplitudes in the DLS. **C-D)** Male and **E-F)** female HFS-LTD time course and 91-100 summary from DLS. Data ± SEM. N=3-10 mice

**Figure S20**

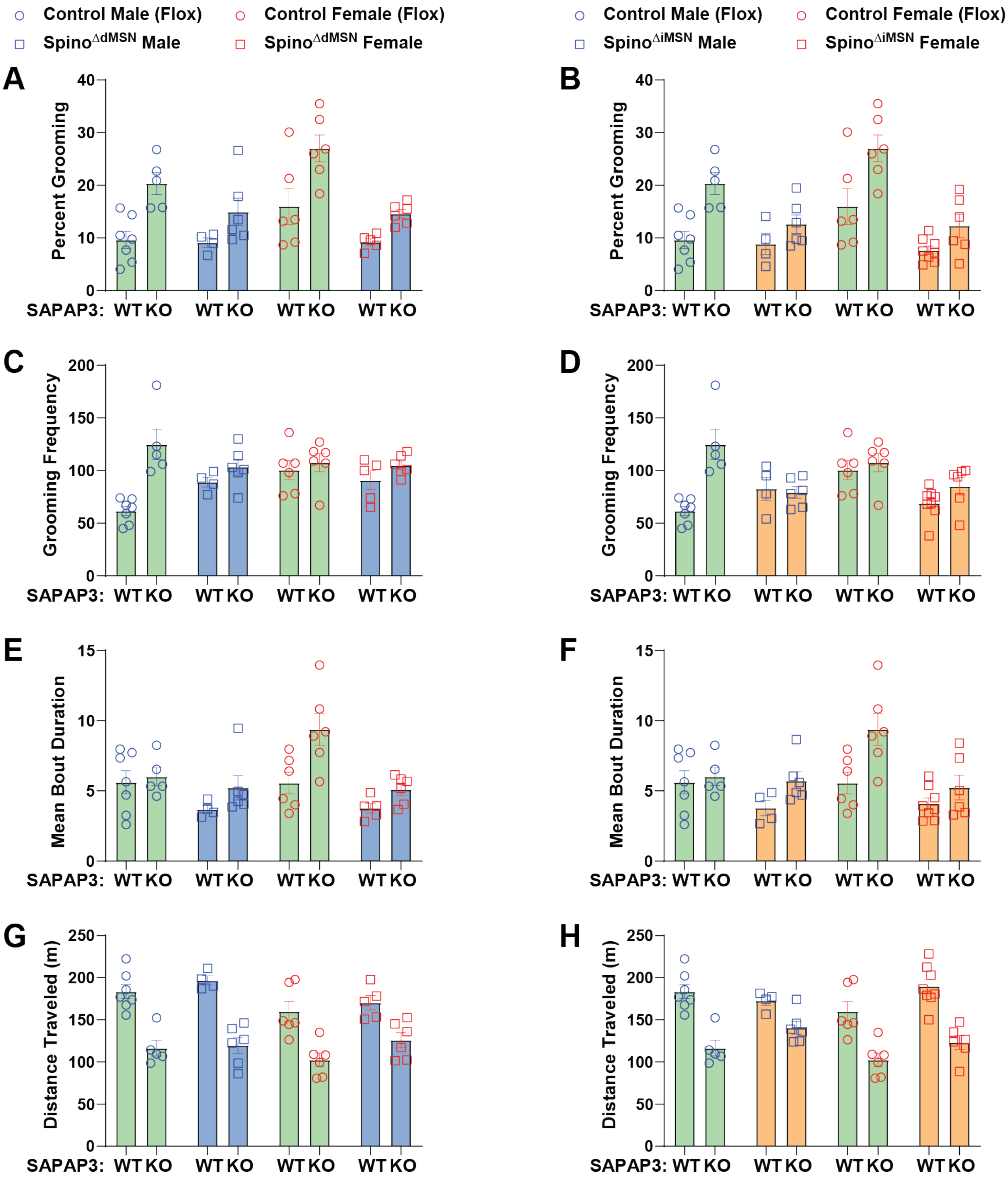


**Figure S20: SAPAP3 WT and KO motor dysfunction by sex.** Male (blue points) and female (red points) SAPAP3 WT-KO/Spino$^{Fl/Fl\text{ or }Fl/+}$ (green bars), SAPAP3 WT-KO/Spino$^{\Delta dMSN}$ (blue bars), and SAPAP3 WT-KO/Spino$^{\Delta iMSN}$ (orange bars) were plotted separately for **A-B.** percent grooming, **C-D.** grooming frequency, **E-F.** mean bout duration, **G-H.** distance traveled. Data ± SEM. N=4-8.

### Figure S21

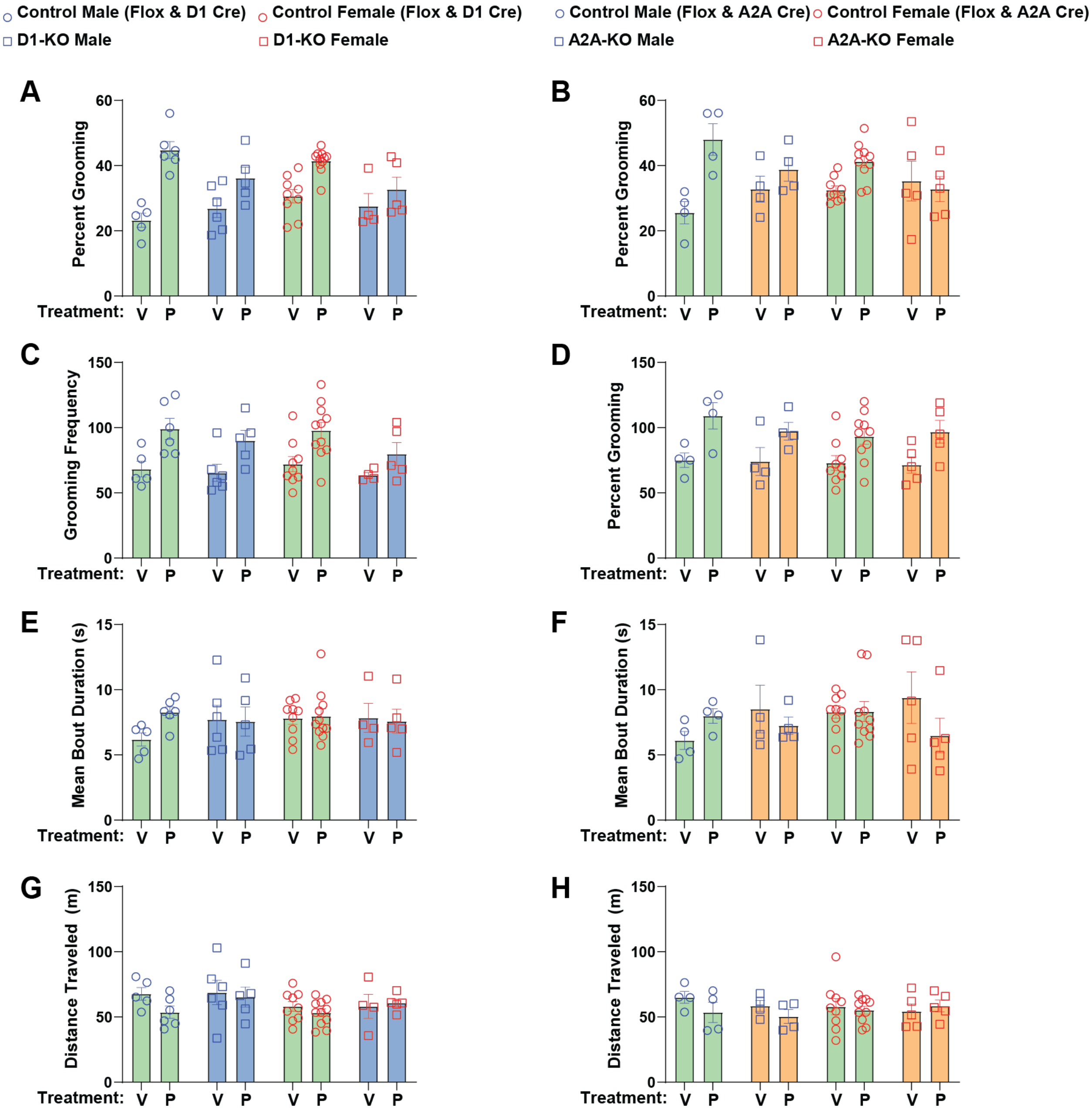


**Figure S21: mGluR5 PAM grooming by sex.** Male (blue points) and female (red points) control ($Spino^{Fl/Fl}$, and $Spino^{+/+}$-D1 or -A2A Cre) (green bars), $Spino^{\Delta dMSN}$ (blue bars), and $Spino^{\Delta iMSN}$ (orange bars) were plotted separately for **A-B.** percent grooming, **C-D.** grooming frequency, **E-F.** mean bout duration, **G-H.** distance traveled. Data ± SEM. N=4-11.

**Figure S22**

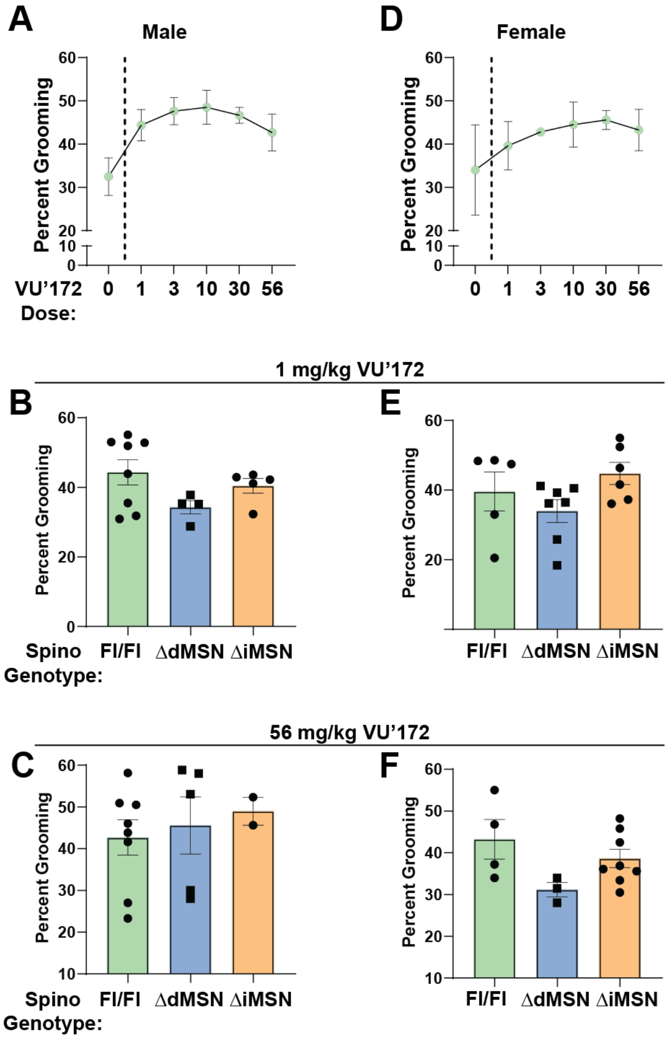


**Figure S22:** Percent grooming data from control dose response, 1 mg/kg VU'172 treatment, and 56 mg/kg VU'172 treatment was plotted separately for **A-C)** males and **D-F)** females. Data ± SEM. N=1-9.

### Supplemental Table Legends

| mGluR5 Peptide | Amino Acid | Abundance Ratio ($Log_2$) |
|---|---|---|
| SSSLVNLWK | Ser860 | 0.40 |
| SPSPISTLSHLAGSAGR | Ser1016 | 0.12 |
| RGSSGETLR | Ser870 | 0.36 |
| SAFTTSTVVR | Ser839 | 0.07 |

**Table S1: Loss of spinophilin increased mGluR5 phosphorylation at mGluR5 Ser860 and Ser870.** Sequential striatal mGluR5 IPs from one spino$^{+/+}$ and one spino$^{-/-}$ mouse were electrophoresed using an SDS-PAGE gel combined with a total protein Imperial Blue stain. mGluR5 bands at ~150kDa were excised from gel for GelC-MS analysis, which identified mGluR5 phosphorylation at Ser839, Ser860, Ser870, and Ser1016**.** Loss of spinophilin increased the abundance ($log_2$-fold change > 0.2) of mGluR5 Ser860 and Ser870.

**Table S2: Protein summary of TMT-LC/MS run 1.** Complete quantitative proteomics tables used to generate protein-protein interaction graphs and gene ontology analysis in string-DB.

**Table S3: Gene ontology summary for top 35 decreased mGluR5 interactions.** Table listing all significant GO terms associated with decreased mGluR5 interactions.

**Table S4: Gene ontology summary for top 35 increased mGluR5 interactions.** Table listing all significant GO terms associated with increased mGluR5 interactions.

**Table S5: Protein summary of TMT-LC/MS run 2.** Complete quantitative proteomics tables used as a targeted validation of TMT-LC/MS run 1.

**Table S6: Statistical Analyses.** All t-, F-, and r-statistics from analyses broken down by experiment and dependent variables.
